# Rice brown spot resistance gene *bsr1* also confers resistance to bacterial blight by suppressing sucrose efflux

**DOI:** 10.64898/2026.08.07.743414

**Authors:** Ritsuko Mizobuchi, Aoi Hishida, Hironori Juichi, Rika Michishita, Fukuyo Tanaka, Yoshihiro Wakabayashi, Haruhiko Inoue, Noriyuki Kuya, Nobuhiro Suziki, Masaki Endo, Masafumi Mikami, Satomi Ohashi, Kengo Matsumoto, Yuya Ota, Tomohiro Yamakawa, Daisuke Nakamura, Chikako Tsuiki, Hiroyuki Sato

## Abstract

Brown spot (BS), caused by the fungal pathogen *Bipolaris oryzae*, is a major disease threatening global rice production. However, the genetic basis of host BS resistance remains unclear. Here, we identified *brown spot resistance 1* (*bsr1*), a quantitative trait locus conferring BS resistance, by map-based cloning. We show that *bsr1* encodes a sucrose transporter and that a near-isogenic line carrying *bsr1* (*bsr1*-NIL) in the susceptible Koshihikari genetic background exhibited resistance to BS by suppressing sucrose efflux into the apoplast after pathogen attack. Furthermore, *bsr1*-NIL also showed strain-specific resistance to bacterial blight caused by *Xanthomonas oryzae* pv. *oryzae* through the same mechanism. These findings demonstrate that *bsr1* confers dual resistance to fungal and bacterial diseases by regulating sucrose efflux. Our study identifies a previously unrecognized mechanism underlying resistance to both BS and bacterial blight and highlights *bsr1* as a promising target for breeding disease-resistance rice cultivars. Rice (*Oryza sativa* L.) is a staple food for more than half of the world’s population^1^. Brown spot (BS), caused by the fungus *Bipolaris oryzae*, is one of the most prevalent fungal diseases of rice, and its incidence has increased under global warming^2^. BS infects coleoptiles, leaves, leaf sheaths, panicle branches, glumes, and spikelets, and severe infection can substantially reduce grain yield.

---

To date, the most significant outbreak of BS was associated with the Bengal famine of 1943^3^, during which a 40–90% reduction in rice yield caused by BS is estimated to have contributed to the deaths of more than two million people in Bengal^3^. More recently, BS outbreaks have also been reported in Brazil^4^, East and Southeast Asia^5^, and India^6^, with reported yield losses of up to 52%^7,8^. In Vietnam, recent outbreaks have caused crop losses of up to 90%^9^. According to a recent report by the Intergovernmental Panel on Climate Change, global mean surface temperatures during 2081–2100 are projected to be 1.0–5.7°C higher than those during the pre-industrial period (1850– 1900)^10^. As a consequence, regions with temperatures favorable for the growth of *B. oryzae*, which grows optimally at around 30°C, are expanding toward higher latitudes, and increased risk and disease severity have been predicted in regions such as the USA and South Africa^11^. Although fungicides are still widely used to control BS, they are costly and pose risks to ecological security. Therefore, breeding rice cultivars with resistance to BS is an important strategy for disease control.

Rice cultivars with varying levels of resistance to BS have been identified^12^, and several quantitative trait loci (QTLs) conferring resistance to BS have been reported^13–20^. However, no genes conferring complete resistance to BS have yet been identified^2^. We previously identified a major BS resistance QTL, *qBSfR11*, later renamed *brown spot resistance 1* (*bsr1*), on chromosome 11 through field evaluations of recombinant lines derived from a cross between the BS-resistant landrace Tadukan and the susceptible cultivar Hinohikari^16,17^. On the basis of this discovery, we developed Mienoyume BSL, the world’s first practical rive cultivar with BS resistance, by introducing *bsr1* into the genetic background of the susceptible cultivar Mienoyume^21^. Because Mienoyume BSL showed a 28.8% higher yield than Mienoyume under severe BS conditions, *bsr1* is expected to be an effective resistance gene against BS^21^. Characterizing the mechanisms underlying *bsr1*-mediated natural resistance to BS is therefore important. However, the molecular and biochemical mechanisms by which *bsr1* regulates defense responses against BS remain unclear.

Here, we show that *bsr1* encodes a sucrose transporter and demonstrate that a near-isogenic line containing *bsr1* (*bsr1*-NIL) in the susceptible Koshihikari genetic background exhibits resistance to BS by suppressing sucrose efflux into the apoplast. In many plant species, SWEET (Sugars Will Eventually Be Exported Transporter) genes function as targets of pathogen effector proteins during host–microbe interactions and contribute to disease susceptibility^22–25^. In rice, loss-of-function mutations of *OsSWEET11*, *OsSWEET13*, and *OsSWEET14* confer resistance to bacterial blight caused by *Xanthomonas oryzae* pv. *oryzae* (*Xoo*)^23^. These findings suggest that host sugar transporters can influence susceptibility to bacterial pathogens. We therefore hypothesized that *bsr1* might also confer resistance to bacterial blight. By inoculating *bsr1*-NIL with several *Xoo* strains, we found that *bsr1*-NIL did indeed exhibit strain-specific resistance to bacterial blight. Our findings provide new insights into a common mechanism underlying resistance to both brown spot and bacterial blight in rice.

## Results

### *bsr1* shows broad resistance to brown spot

Previously, we identified a BS-resistant line carrying a 463-kb chromosomal segment encompassing *bsr1* from the BS-resistant rice cultivar Tadukan in the susceptible Koshihikari genetic background^26^. Because this was the smallest donor segment that retained BS resistance, we used that line as the *bsr1*-NIL throughout the present study. Figure 1a summarizes the genotypes of *bsr1*-NIL and its parental cultivars.

**Fig. 1.**
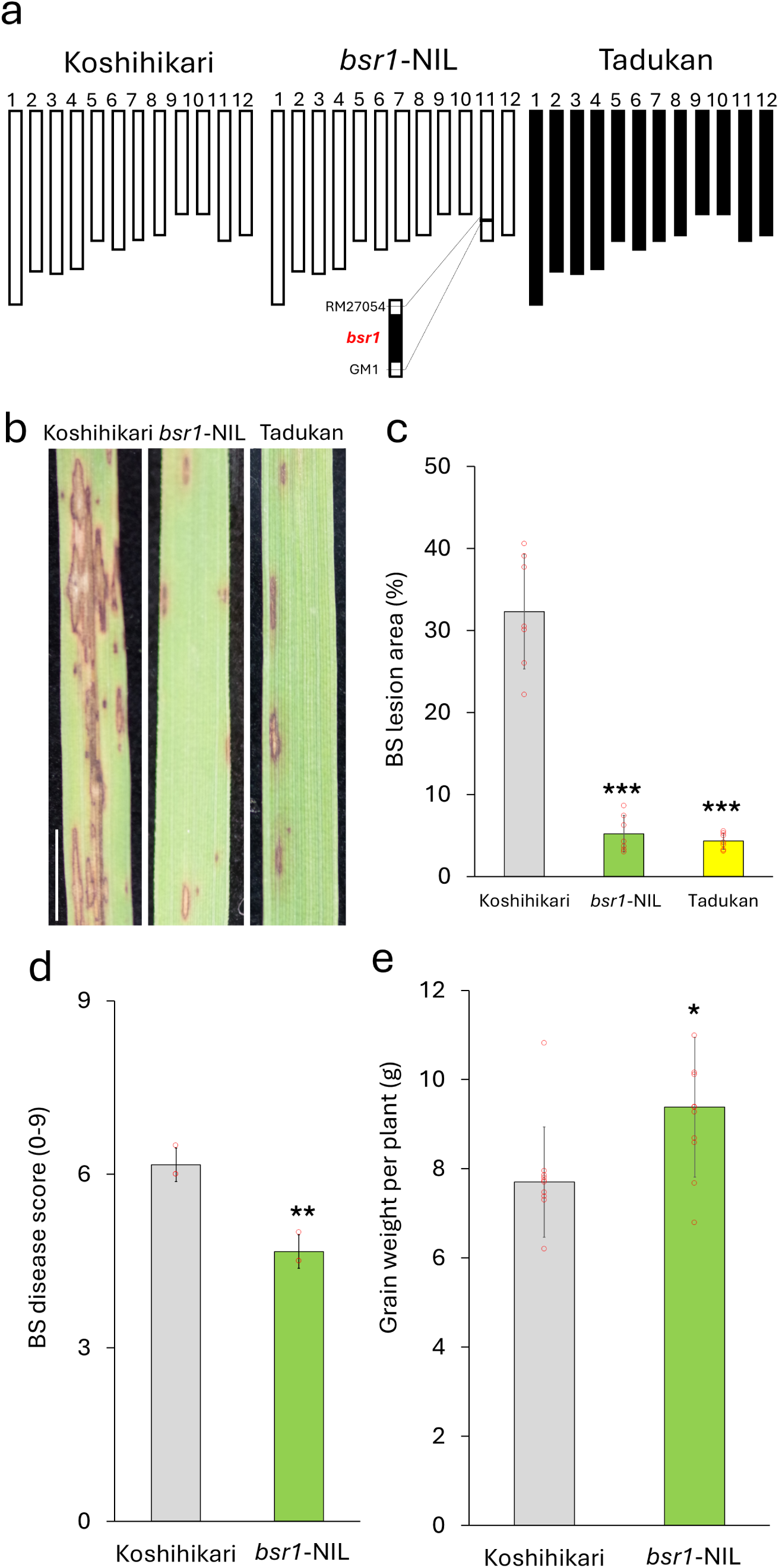
Phenotypic characterization of *brown spot resistance 1* (*bsr1*). **a**, Graphical overview of the genotypes (chromosomes 1–12) of Koshihikari, Tadukan, and the near-isogenic line *bsr1*-NIL. This line carries a 463-kb chromosomal segment encompassing *bsr1* from the brown spot (BS)-resistant rice cultivar Tadukan in the susceptible Koshihikari genetic background; *bsr1* is located between markers RM27054 and GM1 on chromosome 11. White and black rectangles indicate chromosomal regions derived from Koshihikari and Tadukan, respectively. **b,c,** Evaluation of BS resistance in *bsr1*-NIL and its parental cultivars by growth chamber inoculation assay, showing representative BS leaf lesions **(b)** and the quantified lesion area **(c)**. Twenty-seven-day-old plants were inoculated with *Bipolaris oryzae*. Photos were taken 14 days after inoculation, and lesion area was quantified using ImageJ. Scale bar, 5 mm. Data are presented as mean ± s.d. of 7 biological replicates. Asterisks indicate significant differences compared with Koshihikari (Student’s *t*-test, \*\*\**P* < 0.001). **d,e,** Evaluation of BS resistance in Koshihikari and *bsr1*-NIL under field conditions, showing BS disease scores **(d)** and grain weight per plant **(e)**. Disease severity was scored visually at maturity on a scale from 0 (no symptoms) to 9 (severe disease). Data are presented as mean ± s.d. (BS disease score, 11 plants × 3 repeats; grain weight, 10 biological replicates). Asterisks indicate significant differences compared with Koshihikari (Student’s *t*-test, \**P* < 0.05, \*\**P* < 0.01).

To confirm the contribution of the resistance allele of *bsr1* to BS resistance, we compared *bsr1*-NIL with its parental cultivars in a growth chamber inoculation assay. Leaves of *bsr1*-NIL developed significantly smaller BS lesions than those of Koshihikari, and the level of resistance of *bsr1*-NIL was comparable to that of Tadukan (Fig. 1b, c and Supplementary Fig. 1).

Next, to assess the breadth of resistance conferred by *bsr1*, we evaluated the response of *bsr1*-NIL to *B. oryzae* strains collected from different regions of Japan. This line exhibited resistance to all six strains tested, indicating that *bsr1* confers broad-spectrum resistance to *B. oryzae* (Extended Data Fig. 1). In field evaluations, *bsr1*-NIL showed a significantly lower BS disease score than Koshihikari (Fig. 1d) and produced a significantly higher grain weight per plant (Fig. 1e), suggesting that *bsr1* reduces yield loss under severe BS disease pressure.

To determine whether *bsr1* affects agronomic performance, we grew *bsr1*-NIL and Koshihikari in paddy fields where no BS symptoms were observed. Under these conditions, no differences were detected between *bsr1*-NIL and Koshihikari in any of the agronomic traits examined, including grain yield (Extended Data Fig. 2). These results indicate that *bsr1* does not impose a detectable penalty on agronomic performance.

### Map-based cloning of *bsr1*

To clone *bsr1*, we conducted successive rounds of high-resolution mapping and narrowed the candidate interval down to a 31.2-kb region (Fig. 2a and Supplementary Fig. 2). According to the Rice Annotation Project Database (RAP-DB; https://rapdb.dna.affrc.go.jp)^27^, six genes are predicted within this interval (Extended Data Fig. 3a and Supplementary Table 1). Among them, three genes (*Os11g0620400* [*g1*], *Os11g0620450* [*g2*], and *Os11g0621000* [*g6*]) were expressed in inoculated leaves of *bsr1*-NIL (Extended Data Fig. 3b).

**Fig. 2.**
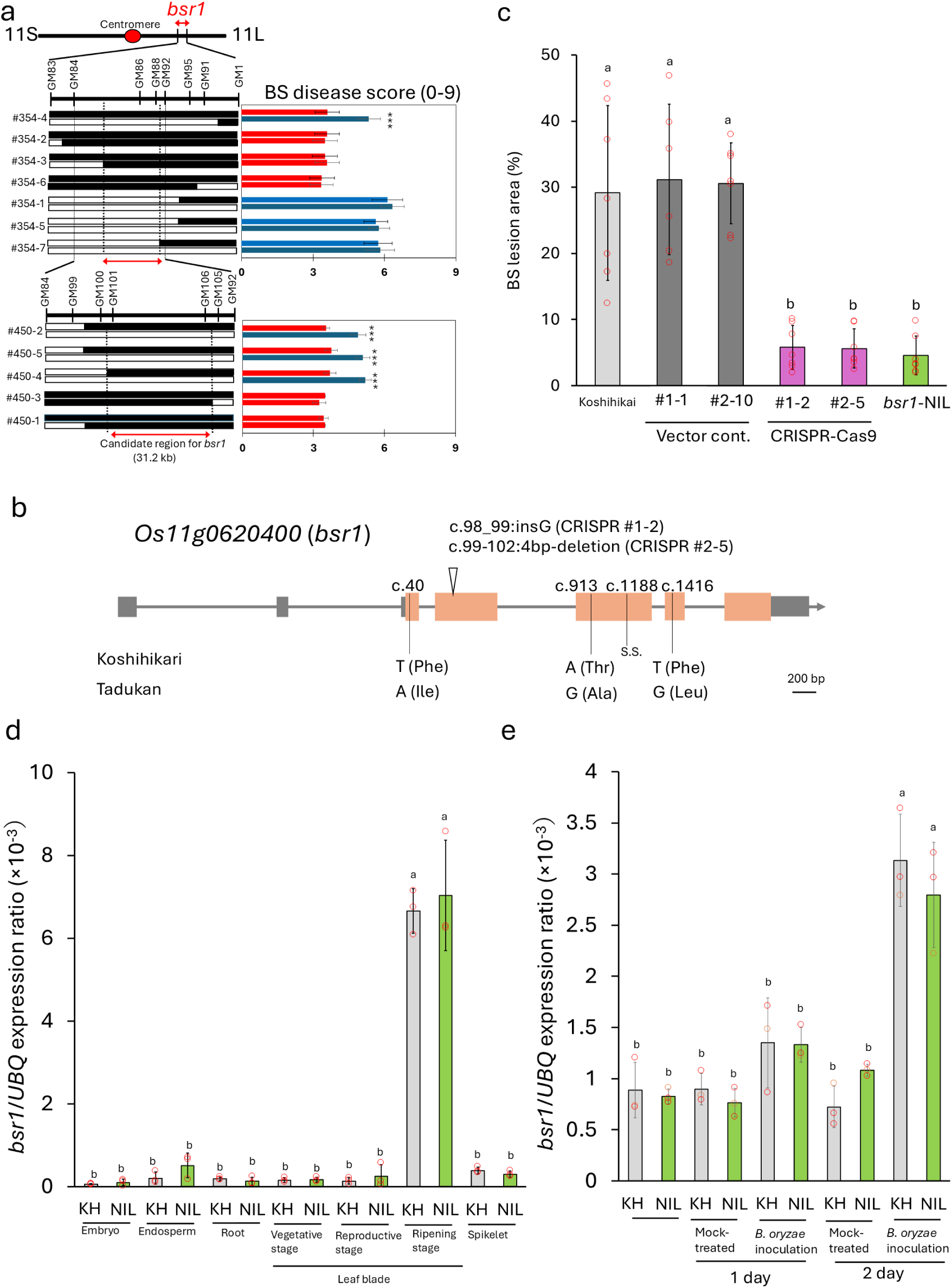
Molecular cloning of *bsr1*. **a**, High-resolution mapping of *bsr1*. In the first round of selection, seven recombinants between markers GM83 and GM1 were selected from 3072 plants. The progeny of each recombinant were evaluated individually for genotype and brown spot (BS) resistance in a BS field test. Based on these results, a second round of mapping was performed by selecting five recombinants between markers GM84 and GM92 from 4032 plants, and their progeny were analyzed individually. White and black rectangles indicate chromosomal regions derived from Koshihikari (susceptible) and Tadukan (resistant), respectively. Marker positions are based on the International Rice Genome Sequencing Project 1.0 pseudomolecules of the Nipponbare genome. Position of the candidate region containing *bsr1* indicated at the bottom is based on the BS field test results summarized on the right. Red and blue rectangles indicate resistant and susceptible phenotypes, respectively. BS disease scores of the two lines in each pair were compared and asterisks indicate significant differences (Student’s *t*-test, \*\*\**P* < 0.001). Data are presented as mean ± s.d. (n = 7–18 plants except #450-3). For line #450-3, only two plants homozygous for the Tadukan allele and six plants homozygous for the Koshihikari allele were available. Therefore, 10 heterozygous plants were included in the analysis to increase the sample size, and no statistically significant difference was detected among the lines. **b,** Gene structure and sequence polymorphisms of *Os11g0620400* (*bsr1*) in Koshihikari and Tadukan. Orange bars represent the coding sequence, and gray bars represent the 5′ and 3′ untranslated regions Arrowhead indicates the CRISPR/Cas9-induced mutation site: a 1-bp insertion (CRIPSR #1 and #2) or a 4-bp deletion (CRIPSR #2–5). s.s., synonymous substitution. **c,** Evaluation of BS resistance in Koshihikari, *bsr1*-NIL, and CRISPR/Cas9 knockout lines of *bsr1*. Plants were inoculated at 27 days of age and BS lesion areas were quantified from scanned leaf images collected 14 days after inoculation using ImageJ. Data are presented as mean ± s.d. (n = 6 or 7 plants). Different letters indicate significant differences (α = 0.05, Tukey’s HSD test). **d,** Relative *bsr1* expression in leaf blades and other tissues of Koshihikari (KH) and *bsr1*-NIL (NIL). Embryo and endosperm samples were collected from mature seeds 5 months after harvest. Root samples were collected 14 days after sowing. Leaf blades from the vegetative, reproductive, and ripening stages were collected 14 days, 77 days, and 105 days after sowing, respectively. Spikelet samples were collected 1 day after flowering. Expression levels of *bsr1* were normalized to that of ubiquitin. Data are presented as mean ± s.d.; n = 3. Different letters indicate significant differences (α = 0.05, Tukey’s HSD test). **e,** Expression of *bsr1* in leaves after *B. oryzae* inoculation under growth chamber conditions in Koshihikari (KH) and *bsr1*-NIL (NIL). Expression levels of *bsr1* were normalized to that of ubiquitin. Data are presented as mean ± s.d.; n = 3. Different letters indicate significant differences (α = 0.05, Tukey’s HSD test).

Comparison of the sequences of these genes between Tadukan and Koshihikari using the TASUKE genome browser in RAP-DB^27^ revealed no differences between the coding sequences of *g2* or *g6*, whereas several single nucleotide polymorphisms (SNPs) were identified in that of *Os11g0620400* (*g1*). These results suggested that *Os11g0620400* was the most likely candidate gene for *bsr1*. *Os11g0620400* is predicted to encode a protein containing a sugar/inositol transporter domain (Supplementary Table 1).

To confirm that *Os11g0620400* is *bsr1*, we generated knockout lines using the clustered regularly interspaced short palindromic repeats (CRISPR)/CRISPR-associated protein 9 (Cas9) system (Fig. 2b). Because *bsr1* is a recessive gene^21^, CRISPR/Cas9 lines were generated in the Koshihikari background. The developed knockout lines displayed significantly smaller BS lesions than Koshihikari plants transformed with the empty vector (Fig. 2c). From these results, we concluded that *Os11g0620400* is *bsr1*.

### Phenotypic and molecular characterization of *bsr1*

To characterize the function of *bsr1*, we compared its expression between Koshihikari and *bsr1*-NIL grown in paddy fields where no BS symptoms were observed. The expression of *bsr1* was highest in leaf blades at the ripening stage, whereas lower expression was detected at the vegetative and reproductive stages, as well as in embryos, endosperms, roots, and spikelets, both in Koshihikari and *bsr1*-NIL (Fig. 2d). The expression patterns were comparable between the two lines. Because *bsr1* is predicted to encode a sucrose transporter, we hypothesized that its high expression during the ripening stage might affect sugar accumulation in leaves. We therefore compared leaf sugar content between Koshihikari and *bsr1*-NIL and found that sucrose levels at the ripening stage were similar in the two lines (Supplementary Fig. 3).

Reverse transcription polymerase chain reaction (RT-PCR) confirmed that *bsr1* was expressed in inoculated leaves of *bsr1*-NIL (Extended Data Fig. 3b). To examine pathogen-induced expression in greater detail, we performed quantitative RT-PCR (qRT-PCR) following *B. oryzae* inoculation. In both Koshihikari and *bsr1*-NIL, *bsr1* expression was specifically induced in leaves after inoculation (Fig. 2e), with similar expression patterns in the two lines. Likewise, expression of the pathogenesis-related (PR) genes *PBZ1* and *PR2* was specifically induced following *B. oryzae* inoculation, and their expression profiles were also similar in Koshihikari and *bsr1*-NIL (Extended Data Fig. 4a, b).

Phylogenetic tree analysis showed that several transporter genes are closely related to *bsr1* (Supplementary Fig. 4). To determine the subcellular localization of BSR1 protein, we introduced enhanced green fluorescent protein (EGFP) fusion constructs containing the full-length Koshihikari or Tadukan allele into rice protoplasts. EGFP fluorescence from the Koshihikari protein completely colocalized with the plasma membrane marker (Fig. 3a–c). In contrast, the Tadukan protein showed a fluorescence similar to that of the EGFP control, indicating loss of specific plasma membrane localization (Fig. 3e–g). The distinct subcellular localization patterns of the Koshihikari and Tadukan BSR1 proteins suggested that sequence differences between the two alleles may influence BSR1 localization.

**Fig. 3.**
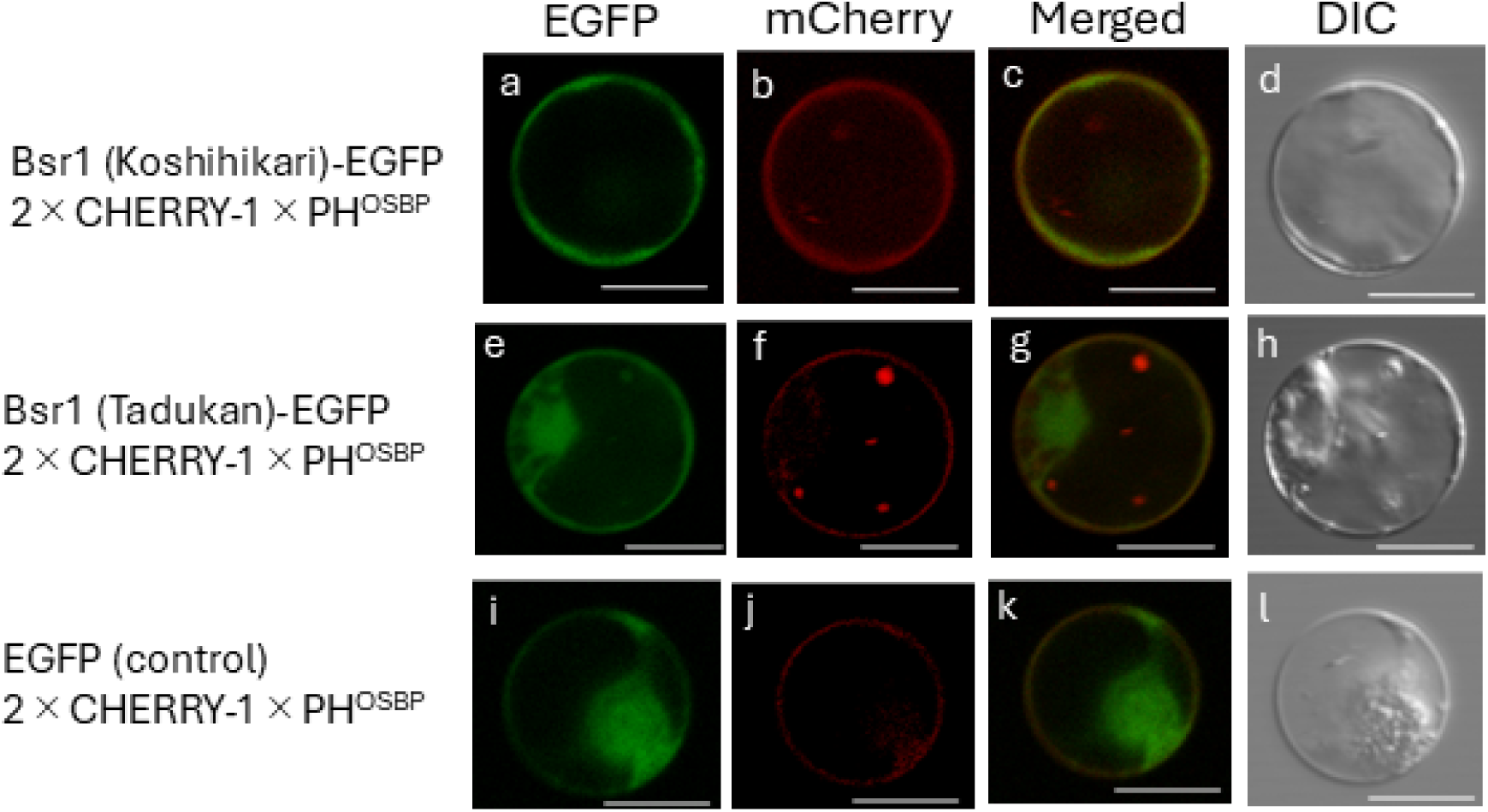
Subcellular localization of Bsr1 in rice protoplasts expressing 2×35Sprom::Bsr1(Koshihikari)-EGFP, 2×35Sprom::Bsr1(Tadukan)-EGFP, or 2×35Sprom::EGFP. **(a,e,i)**. Enhanced green fluorescent protein (EGFP) fluorescence. **b,f,j,** mCherry fluorescence. **c,g,k,** Merged EGFP and mCherry images. **d,h,l,** Differential interference contrast (DIC) images. mCherry fluorescence was induced by the plasma membrane marker UBQ10prom::2×CHERRY-1×PH^OSBP^. Scale bars = 10 μm.

In the *Os11g0620400* coding sequence, one synonymous substitution (SNP1188) and three non-synonymous substitutions (T40A(SNP40), A913G(SNP913) and T1416G(SNP1416)) were identified in Tadukan relative to Koshihikari (Fig. 2b, Supplementary Fig. 5). To evaluate the potential structural effects of these polymorphisms, we predicted the three-dimensional structure of BSR1 using AlphaFold 3^28^ (Supplementary Fig. 6). The predicted structure exhibited the canonical architecture of sugar transporters, comprising 12 transmembrane helices organized into two domains connected by a long intracellular loop between the sixth and seventh transmembrane helices^29^. In this model, Phe14 (Koshihikari-type SNP40) and Phe472 (Koshihikari-type SNP1416) are located within transmembrane helices, whereas Thr305 (Koshihikari-type SNP913) is located within the long intracellular loop.

To identify the SNP responsible for BS resistance, we analyzed the association between BS disease scores obtained in a previous field evaluation^12^ and *Os11g0620400* SNP genotypes in the World Rice Collection and Japanese Rice Collection using sequencing data from TASUKE^30^ (Supplementary Table 2). No significant association was detected for SNP1416 (Supplementary Fig. 7). In contrast, cultivars carrying the Tadukan-type allele at SNP40 or SNP913 showed significantly lower BS disease scores than cultivars carrying the Koshihikari-type allele.

We then examined the distributions of SNP40, SNP913, and SNP1416 in 33 accessions of *Oryza rufipogon*, a wild ancestor of *O. sativa*, using the OryzaGenome database (http://viewer.shigen.info/oryzagenome/mapview/Top.do). All accessions carried the Koshihikari-type nucleotide at SNP40 and SNP1416, whereas four accessions carried the Tadukan-type nucleotide at SNP913 (Supplementary Table 3). These findings suggest that the A913G substitution at SNP913 arose in *O. rufipogon* and was subsequently retained during rice domestication because it confers increased resistance to *B. oryzae*.

### *bsr1* confers resistance to brown spot by suppressing sucrose efflux

Because *bsr1* is predicted to encode a sugar transporter and the Tadukan BSR1 has lost specific plasma membrane localization, we hypothesized that *bsr1* affects sugar distribution in leaves following *B. oryzae* infection. We therefore compared the sugar content of inoculated leaves between Koshihikari and *bsr1*-NIL using three independent approaches: enzymatic sucrose quantification, transfer-based mass spectrometry imaging (MSI), and section-based MSI.

Enzymatic analysis showed that sucrose levels in *B. oryzae*-inoculated leaves of *bsr1*-NIL remained similar to those in mock-inoculated leaves (Fig. 4a). In contrast, sucrose levels in inoculated leaves of Koshihikari were significantly reduced following *B. oryzae* inoculation. Transfer-based MSI yielded the same trend: sucrose signals in *bsr1*-NIL remained comparable to those of mock-inoculated leaves, whereas those in Koshihikari were markedly reduced after inoculation (Fig. 4b, c). Similar results were obtained using the section-based MSI method (Supplementary Fig. 8).

**Fig. 4.**
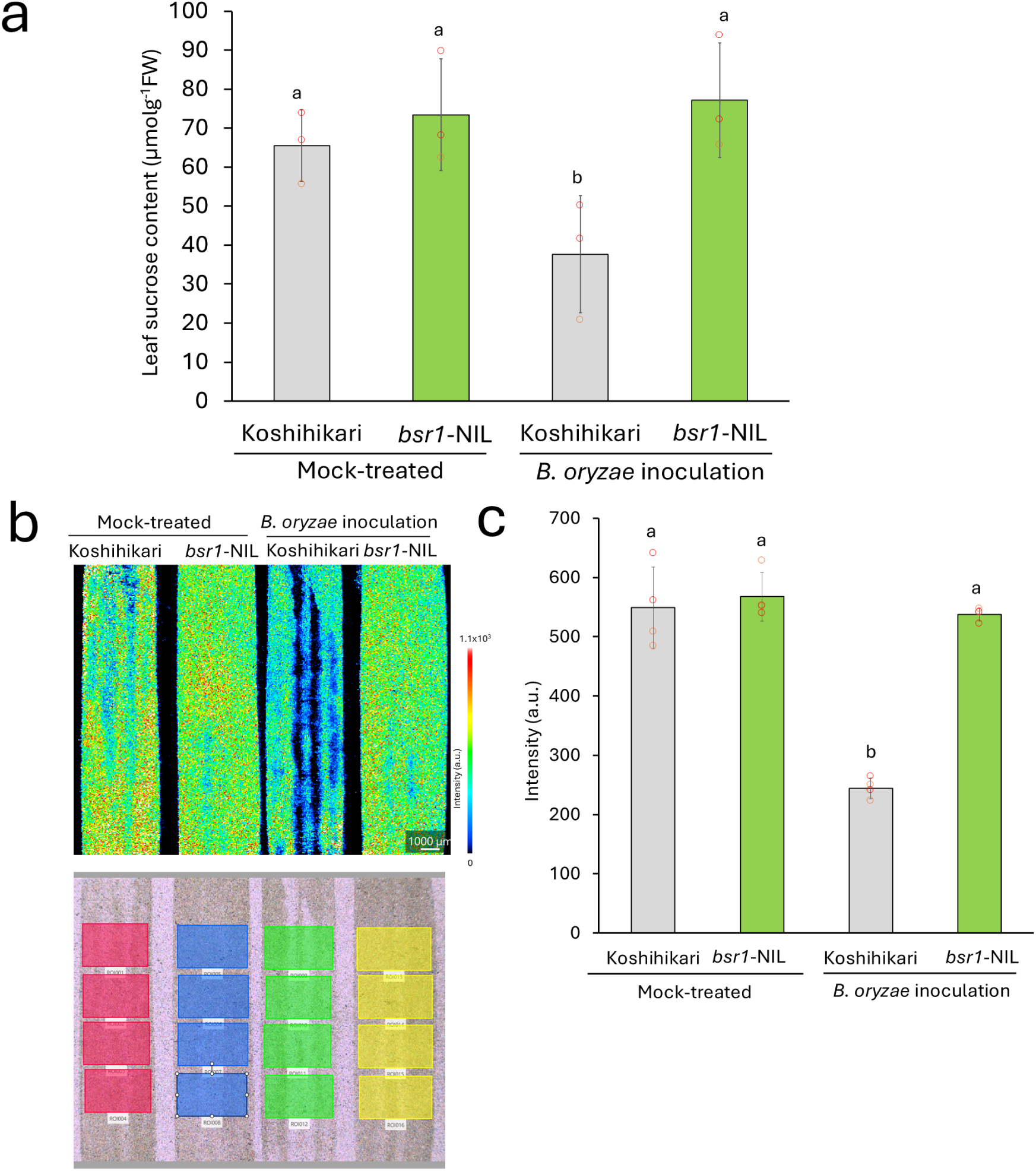
*bsr1* confers brown spot resistance by suppressing sucrose efflux. **a**, Sucrose content of leaves five days after *Bipolaris oryzae* inoculation in Koshihikari and *bsr1*-NIL. Data are presented as mean ± s.d.; n = 3. Different letters indicate significant differences (α = 0.05, Tukey’s HSD test). **b,c,** Sucrose distribution in leaves five days after *B. oryzae* inoculation. **b,** Representative ion images (sucrose, *m*/*z* 381.08) (top) and regions of interest (ROIs) (bottom) ; **c,** quantification of signal intensity. Inoculated leaves were transferred onto platinum-coated porous plates, and sucrose distribution was analyzed by mass spectrometry imaging. Sucrose signal intensity was quantified from regions of ROIs. Data are presented as mean ± s.d. (n = 4 ROIs). Different letters indicate significant differences (α = 0.05, Tukey’s HSD test).

These findings suggest that *bsr1* confers resistance to BS by suppressing pathogen-induced sucrose efflux from host cells into the apoplast.

### *bsr1* also confers strain-specific resistance to bacterial blight

In plants, SWEET proteins play important roles in interactions with pathogens, and rice mutants lacking OsSWEET11, OsSWEET13, or OsSWEET14 exhibit resistance to bacterial blight caused by *Xoo*^23^. We therefore hypothesized that *bsr1* might also confer resistance to bacterial blight in addition to BS.

Inoculation tests with four representative Japanese strains of *Xoo* showed that *bsr1*-NIL was resistant only to strain T7174, (Fig. 5a, b). In Japan, *Xoo* strains are classified into five races based on their virulence patterns on rice cultivars, and T7174 belongs to race I^31^. To determine whether *bsr1*-NIL also showed resistance to other race I strains, we evaluated its response to additional isolates. However, resistance was observed only against T7174 (Supplementary Fig. 9). These results indicate that *bsr1* confers strain-specific resistance to bacterial blight.

**Fig. 5.**
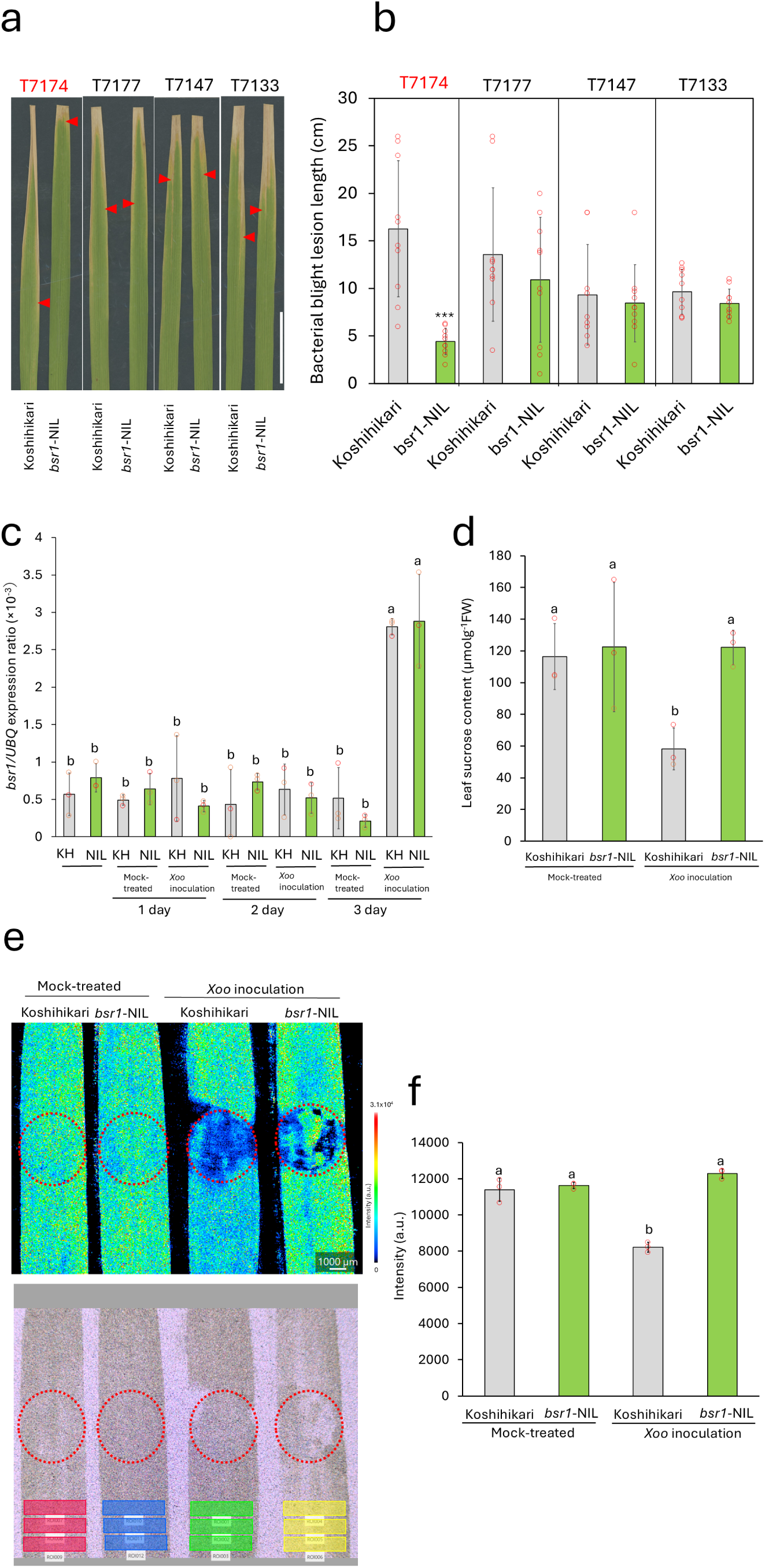
*bsr1* confers bacterial blight resistance by suppressing sucrose efflux into the apoplast. **a,b,** Evaluation of bacterial blight resistance of Koshihikari and *bsr1*-NIL: **(a)** representative lesions; **(b)** quantified lesion lengths. Flag leaves were inoculated with four virulent *Xanthomonas oryzae* pv*. oryzae* (*Xoo*) strains (T7174, T7177, T7147, and T7133) using the clipping method, and lesion lengths were measured 14 days after infection. Data are presented as mean ± s.d. of n = 9 or 10 biological replicates. Asterisks indicate significant differences compared with Koshihikari (Student’s *t*-test, \*\*\**P* < 0.001). **c,** Expression of *bsr1* following inoculation with *Xoo* strain T7174 using the clipping method in Koshihikari (KH) and *bsr1*-NIL (NIL). Expression level of *bsr1* was normalized to that of ubiquitin. Data are presented as mean ± s.d.; n = 3. Different letters indicate significant differences (α = 0.05, Tukey’s HSD test). **d,** Sucrose content of leaves three days after inoculation with *Xoo* strain T7174 by infiltration using a needleless syringe in Koshihikari and *bsr1*-NIL. Data are presented as mean ± s.d.; n = 3. Different letters indicate significant differences (α = 0.05, Tukey’s HSD test). **e,f,** Sucrose distribution in leaves three days after inoculation with *Xoo* strain T7174 by the needleless syringe method. **e,** Representative ion images (sucrose, *m*/*z* 381.08) (top) and regions of interest (ROIs) (bottom) **f,** quantified sucrose signal intensity. Red circles indicate the positions where the bacterial suspension was infiltrated using a needleless syringe. Inoculated leaves were transferred onto a platinum-coated porous plate, and sucrose distribution was analyzed by mass spectrometry imaging. Sucrose intensity was quantified from regions of interest (ROIs) surrounding the inoculation site. Data are presented as mean ± s.d. (n = 3 ROIs). Different letters indicate significant differences (α = 0.05, Tukey’s HSD test).

qRT-PCR showed that *bsr1* expression was induced in the leaves of both Koshihikari and *bsr1*-NIL following inoculation with T7174, with similar expression patterns in the two lines (Fig. 5c). Because *B. oryzae* infection significantly reduced leaf sucrose levels in Koshihikari but not in *bsr1*-NIL (Fig. 4, Supplementary Fig. 8), we hypothesized that *bsr1* similarly affects sucrose dynamics during *Xoo* infection.

Enzymatic sucrose quantification showed that sucrose levels in T7174-inoculated leaves of *bsr1*-NIL remained similar to those in mock-inoculated leaves, whereas sucrose levels in inoculated Koshihikari leaves were significantly reduced (Fig. 5d). Transfer-based MSI produced the same result: sucrose signals remained unchanged in *bsr1*-NIL after inoculation but were markedly reduced in Koshihikari (Fig. 5e, f). These findings suggest that the BS resistance gene *bsr1* also confers resistance to bacterial blight by suppressing pathogen-induced sucrose efflux from host cells into the apoplast.

## Discussion

No resistance gene against *B. oryzae* has been cloned to date. In this study, we identified the gene underlying *bsr1*, a BS resistance QTL, which is predicted to encode a sucrose transporter, and *bsr1*-NIL exhibited resistance to BS by suppressing sucrose efflux into the apoplast. Interestingly, *bsr1*-NIL also showed strain-specific resistance to bacterial blight.

Sugars are essential for plant growth and development. Among the gene families involved in sugar transport, three transporter families play key roles in intercellular sugar transport: MST (monosaccharide transporters), SWEET, and SUT (sucrose transporters)^32^; *bsr1* belongs to the MST family^32^. MST proteins are involved in various aspects of plant growth and development and have been characterized in many plant species, including *Arabidopsis thaliana*, *Vitis vinifera*, *Nicotiana tabacum*, *Solanum lycopersicum*, *Medicago truncatula*, and *Rosa hybrida*^32^. According to Deng *et al.*^32^, MST gene expression patterns suggest that these transporters also play important roles in responses to abiotic stress. However, their roles in biotic stress remain largely unknown. To our knowledge, *bsr1* is the first *MST* gene reported to confer resistance to a pathogen. In contrast to MST genes, SWEET genes have been well characterized as key regulators of plant–pathogen interactions^23–25,33,34^. Several SWEET genes associated with pathogen susceptibility have been identified in many plant species, including rice, *Arabidopsis*, sweet potato, cotton, cassava, and citrus^22,23^. It is therefore tempting to speculate that SWEET proteins serve as host susceptibility factors for a wide range of pathogens. *Xanthomonas* transcription activator-like (TAL) effectors are well known to promote disease by binding to and activating host susceptibility genes^35^. In rice, the expression of several OsSWEET genes is induced by TAL effectors from *Xoo*, and three OsSWEET genes have been associated with susceptibility to *Xoo*^36^. The recessive resistance loci *xa13*, *xa25*, and *xa41* correspond to *OsSWEET11*^37–40^, *OsSWEET13*^41^, and *OsSWEET14*^42–44^, respectively. Although TAL effectors have so far been identified only in *Xanthomonas* species and *Ralstonia solanacearum*^45^, *OsSWEET11* mutants, originally identified through their resistance to *Xoo*, also exhibit resistance to sheath blight caused by *R. solani*^37^.Therefore, it is possible that *B. oryzae* possesses proteins functionally similar to TAL effectors, which influence the expression of *bsr1*.

*bsr1*-NIL exhibited broad-spectrum resistance to *B. oryzae,* showing resistance to all strains tested (Extended Data Fig. 1). In contrast, *bsr1*-NIL was resistant only to *Xoo* strain T7174 (Fig. 5a, b and Supplementary Fig. 9). T7174 is predicted to encode 17 TAL effectors based on its genome sequence^45,46^. Although the *bsr1* promoter does not appear to contain a perfect match to previously reported TAL effector–binding elements^35,44^, it remains possible that one of the TAL effectors from T7174 recognizes the *bsr1* promoter. In contrast to *Xoo*, physiological races have not been reported for *B. oryzae*. Therefore, it is possible that the *B. oryzae* strains examined in this study possess a common TAL effector that targets *bsr1*.

In 1985, wheat blast caused by *Magnaporthe oryzae* (which also causes rice blast) emerged in Brazil, and during its subsequent evolution, the powdery mildew resistance gene *Pm4* was found to confer dual resistance to both wheat mildew and wheat blast^47,48^. How *bsr1* confers resistance to both BS and bacterial blight remains unknown. Following the initial intracellular penetration of epidermal cells during compatible interactions with *B. oryzae*, fungal hyphae grow through the intercellular spaces^9^. This observation suggests that *bsr1* may confer resistance to both BS and bacterial blight through a common mechanism by suppressing pathogen-induced sucrose efflux from cells into the apoplast. Further studies will be required to clarify the role of *bsr1* in plant– pathogen interactions.

The recessive bacterial blight resistance gene *xa13*, which corresponds to *OsSWEET11*, does not result in the typical hypersensitive resistance response in rice^38^. Similarly, *bsr1*, which we identified here as a gene conferring resistance to both BS and bacterial blight, suppresses lesion expansion also without inducing a typical hypersensitive resistance response (Figs. 1b and 5a). These observations suggest that *bsr1*-mediated resistance is mechanistically distinct from immunity mediated by genes encoding nucleotide-binding leucine-rich-repeat proteins ^49^. Ethylene has also been implicated in susceptibility to BS and bacterial blight^50–52^. Whether *bsr1* functions through ethylene signaling remains unknown and warrants further investigation.

EGFP fluorescence of the Koshihikari BSR1 protein completely colocalized with the plasma membrane marker (Fig. 3a–c), whereas plasma membrane localization was largely lost in the Tadukan BSR1 protein (Fig. 3e–g). These observations suggest that sequence differences between the two alleles alter the subcellular localization of BSR1. Three non-synonymous substitutions (SNP40, SNP913, and SNP1416) distinguish the Koshihikari and Tadukan alleles (Fig. 2b, Supplementary Fig. 5), and SNP40 and SNP913 were significantly associated with BS resistance (Supplementary Fig. 7). Comparison of these polymorphisms among 33 *O. rufipogon* accessions revealed that all of them carried the Koshihikari-type nucleotide at SNP40 and SNP1416, whereas four accessions carried the Tadukan-type nucleotide at SNP913 (Supplementary Table 3). These accessions originated from Southeast Asian countries, including the Philippines, Indonesia, and Bangladesh. These findings are consistent with the hypothesis that the A913G substitution at SNP913 originated in *O. rufipogon* and was subsequently retained during rice domestication because of its contribution to BS resistance.

*bsr1* was highly expressed in leaf blades at the ripening stage, whereas lower expression levels were detected in other tissues (Fig. 2d). Under non-infected conditions, sucrose levels in leaves at the ripening stage were similar between Koshihikari and *bsr1*-NIL (Supplementary Fig. 3). Mienoyume BSL, which we developed as the world’s first practical cultivar with BS resistance by introducing *bsr1* into the susceptible cultivar Mienoyume, exhibited agronomic performance comparable to that of Mienoyume under BS-free conditions^21^. Similarly, *bsr1*-NIL showed no detectable differences from Koshihikari in agronomic traits under BS-free conditions (Extended Data Fig. 2). Together, these findings indicate that the resistance *bsr1* allele does not impose a detectable agronomic penalty.

*bsr1*-NIL exhibited broad-spectrum resistance to *B. oryzae*, being resistant to all strains tested (Extended Data Fig. 1). In contrast, resistance to *Xoo* was strain specific. Although further studies are needed to clarify the mechanisms underlying resistance to *B. oryzae* and *Xoo*, dual-resistance genes are rare and highly valuable for rice breeding. Our findings provide new insights into the natural regulatory mechanisms underlying resistance to both brown spot and bacterial blight.

## Methods

### Plant materials

Tadukan is a traditional *indica* rice landrace originating from the Philippines that is resistant to BS caused by *B. oryzae.* Koshihikari is a modern lowland *japonica* rice cultivar developed in Japan that is susceptible to BS. To characterize *bsr1*, we developed a NIL homozygous for the Tadukan allele of *bsr1* by crossing Wa3663 with Koshihikari (Supplementary Fig. 2). Wa3663 was previously selected as a BS-resistant line and carries a 1.4-Mbp chromosomal segment from Tadukan on chromosome 11 (between markers IDR2641 and RM27163)^17,21^. The resulting *bsr1*-NIL contains an approximately 463-kb Tadukan-derived segment on the long arm of chromosome 11.

### Measurement of agronomic traits

Agronomic traits were evaluated in a paddy field of the NARO Institute of Crop Science (Tsukuba, Ibaraki; 36.03°N, 140.10°E). Thirty-day-old seedlings were transplanted at a density of one seedling per hill. Each cultivar was planted in a single row of 12 hills with 18 cm between hills and 30 cm between rows. Nitrogen fertilizer was applied at 5.6 g m^−2^.

### Assessment of BS resistance by field test

For mapping *bsr1*, field evaluations of BS resistance were conducted at the Iga Agricultural Research Laboratory of the Mie Prefecture Agricultural Research Institute (Iga, Mie, Japan; 34.70°N, 136.13°E). Seedlings were transplanted in late May in three replicates, with 11 plants per row at a spacing of 30 × 15 cm. Slow-release nitrogen fertilizer was applied at 7.5 g N m^−2^ at transplanting. As spreader plants, seedlings of a susceptible cultivar were inoculated with *B. oryzae* before transplanting and then planted around, but not within, the experimental rows. Because BS symptoms develop gradually, resistance was assessed at maturity using a scale from 0 (no symptoms) to 9 (severe disease)^53^.

### Assessment of BS resistance by growth chamber inoculation test

*Bipolaris oryzae* strains (Supplementary Table 4) were obtained from the NARO Genebank culture collection and cultured on potato dextrose agar medium containing 5% agar at 28°C for 14 days in the dark. Conidiophore formation was then induced by irradiation with black light for 4 days. Seeds of the test plants were sown in sterilized soil (Bonsol No. 2, Sumitomo) in 1.5 × 1.5 × 2 cm cell trays and grown in a growth chamber at 28°C under a 14-h photoperiod. Three weeks after sowing, plants were inoculated with 50 mL of conidial suspension (8 × 10^4^ conidia mL^−1^), maintained at 28°C and 100% humidity for 20 h, and then returned to the growth chamber under the same conditions. Seven days after inoculation, leaves were detached and immersed in 70% ethanol to remove chlorophyll. Leaf images were scanned using an Epson GT-X820 scanner (Seiko Epson), and lesion area was quantified using ImageJ (version 1.53).

### Inoculation with bacterial blight strains

*Xoo* strains obtained from NARO Genebank (Supplementary Table 4) were cultured on potato sucrose agar medium containing 2% agar at 28°C for 4 days and suspended in sterile water. Bacterial suspensions were adjusted to an optical density of OD_660_ = 0.25 before use. To evaluate resistance to bacterial blight, flag leaves of 15-week-old plants grown in an experimental paddy field were inoculated using the clipping method described previously^54^, and lesion lengths were measured 14 days after inoculation.

For MSI, seeds of the test plants were sown in sterilized soil (Bonsol No. 2) in 3.5 × 3.5 × 3.5 cm cell trays and grown in a growth chamber at 28°C under a 14-h photoperiod. Three weeks after sowing, fully expanded leaves were inoculated with *Xoo* strain T7174 by infiltrating the bacterial suspension with a needleless syringe. Three days after inoculation, the leaves were transferred to a platinum-coated porous plate (Poropare, Hamamatsu Photonics)^55^ and subjected to MSI analysis.

### Mapping of *bsr1*

We previously mapped *bsr1* between the simple sequence repeat markers RM27054 and RM27163 on chromosome 11^21^ and selected Wa3663 as a BS-resistant line in the Koshihikari genetic background. For high-resolution mapping of *bsr1*, recombinant homozygous lines or plants were selected in three successive rounds. First, we generated 1344 F_2_ plants by crossing Wa3663 and Koshihikari, and identified 40 plants carrying recombination events within the *bsr1* region (Supplementary Fig. 2). These plants were self-pollinated, and their homozygous progeny were evaluated for BS resistance in field tests. Then, among 3072 plants, we selected 7 plants carrying recombination events within the narrowed *bsr1* interval between markers GM83 and GM1. Finally, among 4032 plants, we selected 5 plants carrying recombination events within the further narrowed interval between markers GM84 and GM92. In the second and third rounds, self-pollinated progeny (approx. 17–30 plants per recombinant line) were transplanted to the BS test field, and each plant was genotyped and evaluated for BS disease severity (Fig. 2a). Total DNA was extracted from fresh leaves as described previously^56^ and used for PCR amplification and electrophoresis as described previously^57^; the primer pairs used are listed in Supplementary Table 5.

### Production of transgenic plants

CRISPR/Cas9 target sites in *bsr1* were designed using CRISPR-P 2.0 (https://cbi.hzau.edu.cn/CRISPR2/), and the vectors were constructed using a previously published method^58^. Guide RNA expression cassettes were cloned into the pZDgRNA binary vector using the AscI and PacI restriction sites. The primers used are listed in Supplementary Table 5. The constructs were introduced into *Agrobacterium tumefaciens* strain EHA101 by electroporation. *Agrobacterium*-mediated rice transformation was then performed as described previously^59,60^. Sequencing of the CRISPR/Cas9 cleavage site in approximately 96 T_0_ seedings identified two types of mutations: a G insertion at c.98_99 and a 4-bp deletion at C.99_102. Homozygous T_1_ plants were identified by sequencing. From progenies of T_1_ seedlings, control plants were selected using the hygromycin phosphotransferase gene and confirmed by sequencing of the CRISPR/Cas9 cleavage site.

### Gene expression analysis

Total RNA was extracted using a RNA Suisui kit (Rizo), and first-strand cDNA was synthesized using SuperScript II Reverse Transcriptase (Invitrogen). qRT-PCR was performed in triplicate using a QuantStudio 6 Pro real-time PCR system (Thermo Fisher Scientific) with Thunderbird SYBR qPCR Mix (QPS-201, Toyobo). For analysis of *PBZ1* and *PR2* expression, TaqMan probes were used with qPCR Master Mix (RT-QP2X-03, NipponGene). Relative expression levels were normalized to that of ubiquitin. The primers used for gene expression analysis are listed in Supplementary Table 5.

### Sequence alignment and phylogenetic analysis

For *bsr1* mapping and confirmation of the transgenic plants, genomic sequences of *bsr1* were amplified by PCR using the primer pairs listed in Supplementary Table 5. PCR products were sequenced by using a BigDye Terminator v.3.1 Cycle Sequencing Kit (Life Technologies). Sequence information for the cultivars analyzed was obtained from the TASUKE genome browser of the RAP-DB (https://rapdb.dna.affrc.go.jp). Homologous proteins were identified by BLAST searches using the BSR1 amino acid sequence as the query. A phylogenetic tree was constructed using Genetyx ver.16 (Genetyx) with the neighbor-joining method and 1000 bootstrap replicates.

### Subcellular localization analysis

The open reading frame sequences of the Tadukan and Koshihikari *bsr1* alleles were amplified by PCR from cDNA using the primers listed in Supplementary Table 5. The amplified fragments were subcloned into pENTER4 (Invitrogen) using an In-Fusion Directional Cloning Kit (Takara). After sequence verification, the plasmids were recombined into the destination vector pSAT6-DEST-EGFP-N1 using Gateway LR Clonase (Invitrogen). The resulting constructs—2 × 35Sprom::bsr1(Tadukan)-EGFP, 2 × 35Sprom::bsr1(Koshihikari)-EGFP, 2 × 35Sprom::EGFP, and the plasma membrane marker UBQ10prom::2×CHERRY-1×PH^OSBP^ ^61^—were transfected into protoplasts prepared from young rice seedlings using a previously described method with a slight modification^62,63^. Stems from 1-week-old rice seedlings were cut into 1-cm segments with a razor blade and digested in cellulase solution containing 10 mM β-mercaptoethanol. Protoplasts were prepared as described previously^63^, and 40 μL of protoplast suspension (ca. 1000 cells) was transfected with 1 μg of plasmid DNA. After overnight incubation, EGFP and mCherry fluorescence was observed using a LSM 710 confocal laser scanning microscope (Carl Zeiss), according to the manufacturer’s protocol.

### Sample preparation for MSI

Two sample preparation methods were used for MSI analysis: a transfer-based method and a section-based method.

*Transfer-based method*: Rice leaves inoculated with *B. oryzae* or *Xoo* were transferred onto a platinum-coated porous plate (Poropare) without sectioning or matrix application, as described previously^55^. Leaf tissues were placed on the plate and covered with paraffin film to prevent movement. The tissues were then repeatedly and uniformly pressed onto the plate surface using a cylindrical rod to transfer molecular components by direct imprinting until a clear leaf morphology became visible on the plate, while avoiding visible attachment of tissue fragments.

*Section-based method*: Rice leaves inoculated with *B. oryzae* were embedded in optimum cutting temperature compound (Sakura Finetek Japan) and rapidly frozen in a dry ice–ethanol bath. Frozen samples were sectioned at a thickness of 10 μm using a cryostat (HM525 NX, Thermo Fisher Scientific) and mounted onto indium tin oxide– coated glass slides (Matsunami Glass Ind.; surface resistivity, 100 Ω/sq). α-Cyano-4-hydroxycinnamic acid was uniformly applied as the matrix using an automated matrix deposition system (iMLayer, Shimadzu) to a thickness of 0.7 μm prior to MSI analysis.

### MSI measurements

MSI measurements were performed using an iMScope QT instrument (Shimadzu), which integrates an optical microscope with an atmospheric-pressure matrix-assisted laser desorption/ionization ion source coupled to a quadrupole time-of-flight mass spectrometer. Data were acquired in positive ion mode over an *m*/*z* range of 300–500. The acquisition parameters were as follows: laser repetition rate, 1000 Hz; spatial resolution, approx. 50 μm; laser energy, 70% of the maximum output; and 50 laser shots per pixel. The heat block and desolvation line temperatures were maintained at 450°C and 250°C, respectively. Unless otherwise specified, all measurements were performed under these conditions.

### Data acquisition and processing

MSI data were acquired using Imaging MS Solution software (Shimadzu). Ion images were generated using IMAGEREVEAL MS software (Shimadzu) based on the signal intensity at *m*/*z* 381.08 assigned to the potassium adduct of sucrose ([M+K]^+^), with a mass tolerance of ±0.01 Da. Region-of-interest analysis was performed using the same software, and signal intensities were extracted for quantitative comparison.

### Enzymatic sucrose quantification

About 5 mg fresh weight of leaf tissue was frozen in liquid nitrogen, ground to a fine powder, and homogenized in 1 mL of sterilized water. The homogenates were incubated in a water bath at 60°C for 20 min and then used for sucrose quantification with a Sucrose/D-Glucose Assay Kit (Megazyme).

### Prediction of protein structure of BSR1

The three-dimensional structure of BSR1 was predicted using AlphaFold 3^28^.

### Statistical analyses

Statistical analyses were performed using JMP v.14.0.0. (SAS Institute).

### Reporting summary

Further information on the research design is available in the Nature Portfolio Reporting Summary linked to this article.

## Data availability

The datasets generated during this study are available from the DNA Data Bank of Japan (DDBJ; https://www.ddbj.nig.ac.jp). The DDBJ accession numbers for *bsr1* from ‘Tadukan’ and ‘Koshihikari’ are LC944043 and LC944042, respectively. All plasmids, plant lines, and fungal strains used or generated in this work are available from the corresponding author upon reasonable request. Source data are provided with this paper. Additional data supporting the findings of this study are available from the corresponding author upon reasonable request.

## Acknowledgements

We are thankful to Dr. Y. Jaillais (ENS Lyon) for providing the UBQ10 prom::2×CHERRY-1×PH^OSBP^ vector, S. Sakamoto (AIST) for his advice on the protoplast experiments, and T. Kawai (NARO) for his valuable support with the microscopy experiments. We thank the NARO technical support staff and the field managers of the Mie Prefecture Agricultural Research Center. This work was supported by JSPS KAKENHI Grant 24K01734 and by a grant from the Ministry of Agriculture, Forestry and Fisheries of Japan (Research Program on Development of Improved Crop Varieties for Food Security).

## Author Contributions

R.Mizobuchi and H.S. designed the experiments and managed the projects. H.J., R.Michishita, F.T., and Y.W. performed mass spectrometry imaging analysis. H.I., N.K., and A.H. performed subcellular localization analysis. N.S. analyzed the AlphaFold-predicted structure of BSR1. M.E. and M.M. performed with transformation experiments. S.O., K.M., Y.O., T.Y., and D.N. performed the brown spot field test. C.T. performed screening of germplasms and their molecular analyses. All authors read and approved of the final manuscript.

---

**Extended Data Fig. 1.**
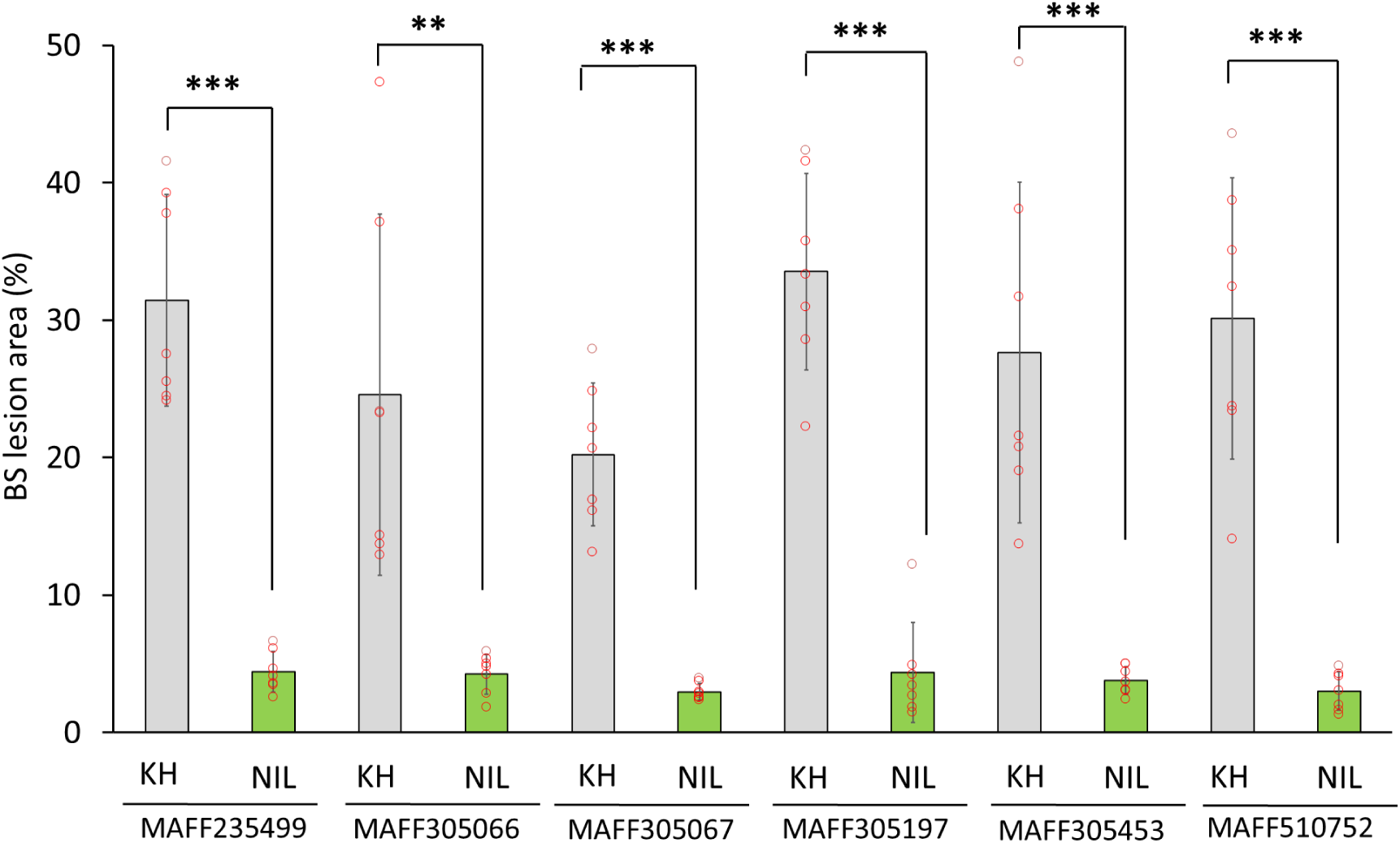
Brown spot lesion area in 27-day-old Koshihikari (KH) and *bsr1*-NIL (NIL) plants inoculated with six *Bipolaris oryzae* strains under growth chamber conditions. The plants were inoculated, and lesion areas were quantified from scanned leaf images collected 14 days after inoculation using ImageJ. Data are presented as mean ± s.d. of 7 biological replicates. Asterisks indicate significant differences compared with Koshihikari (Student’s *t*-test, \*\**P* < 0.01, \*\*\**P* < 0.001).

**Extended Data Fig. 2.**
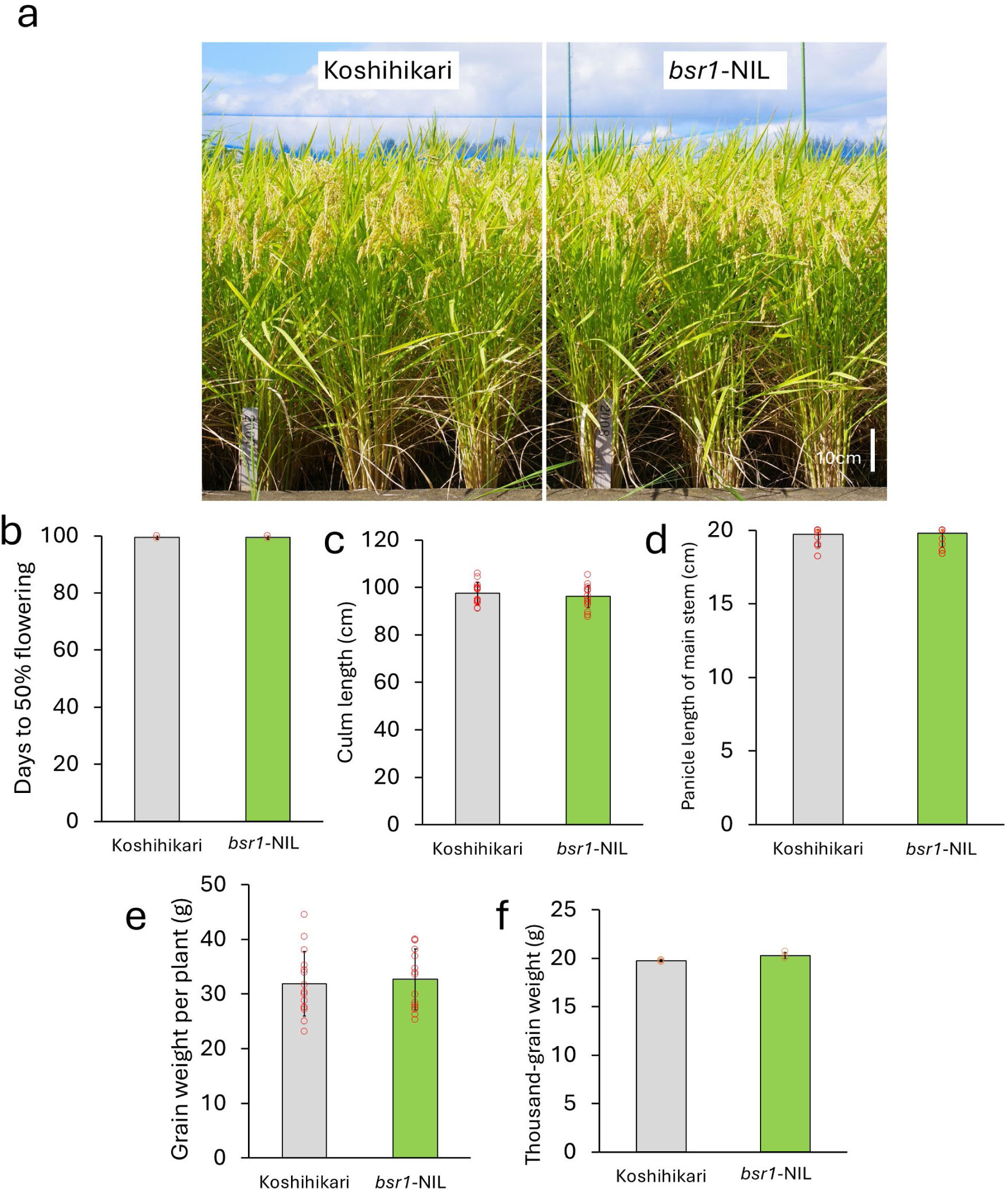
Comparison of agronomic performance between Koshihikari and *bsr1*-NIL. **a**, Representative photographs of Koshihikari and *bsr1*-NIL plants grown in a paddy field. **b,** Days to 50% flowering. Data are presented as mean ± s.d.; n = 2 blocks (12 plants per block). **c–e,** Yield-related traits measured at harvest, including culm length **(c)**, panicle length of the main stem **(d)**, and grain weight per plant **(e)**. Data are presented as mean ± s.d. (n = 15 plants). **f,** Thousand-grain weight. Each group consisted of 4 plants. A 25-g grain sample was collected from each group, and the number of grains was counted. Data are presented as mean ± s.d. (n = 3 groups).

**Extended Data Fig. 3.**
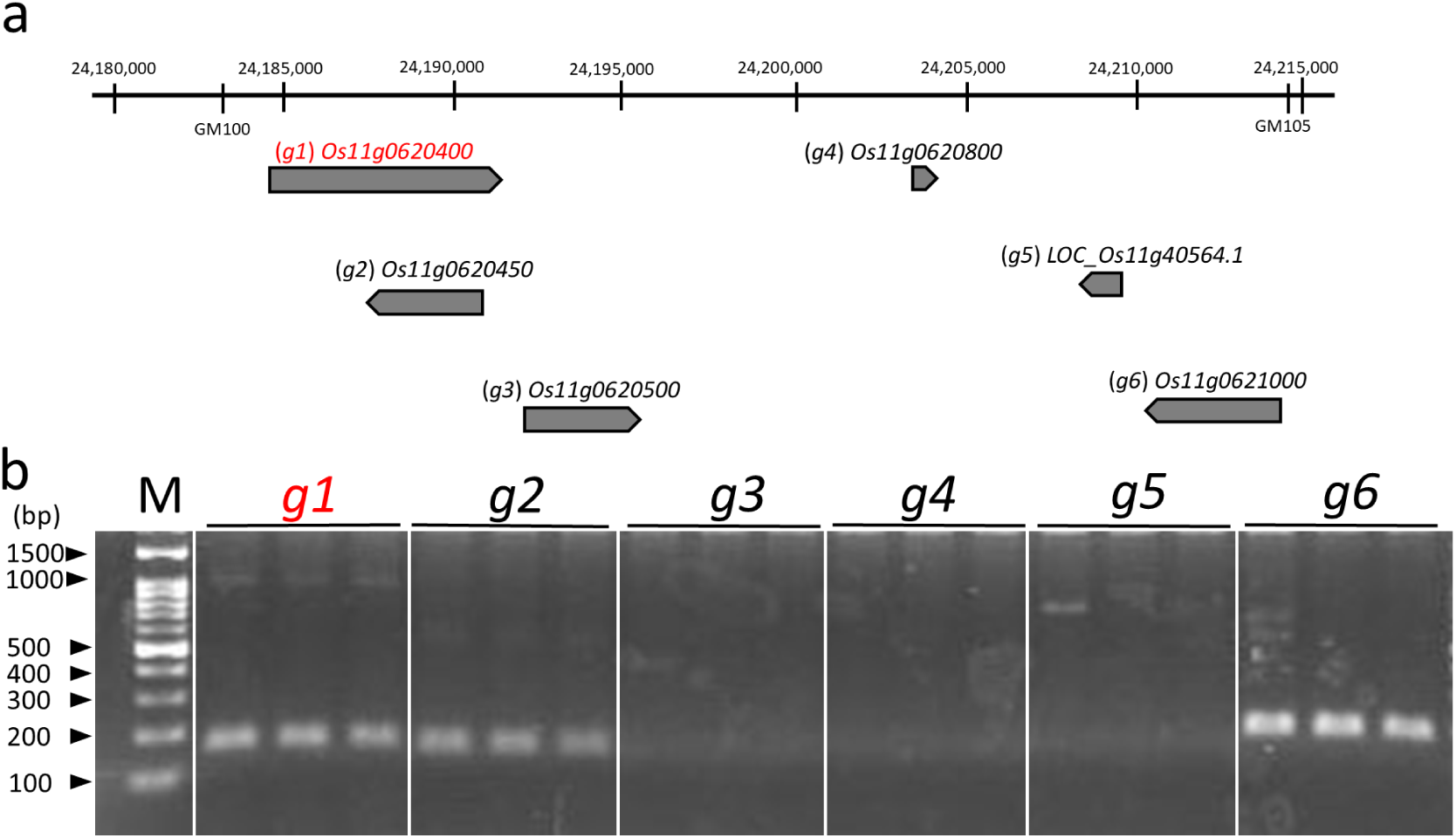
Delimitation of the *bsr1* locus to a 31.2-kb region of chromosome 11. **a**, Predicted genes within the interval flanked by markers GM100 and GM105. Markers and genome information were obtained according to the Nipponbare reference genome in the Rice Annotation Project Database. **b,** Expression of the predicted genes in the leaves of *bsr1*-NIL after inoculation of 27-day-old plants with *Bipolaris oryzae*, as determined by reverse transcription polymerase chain reaction.

**Extended Data Fig. 4.**
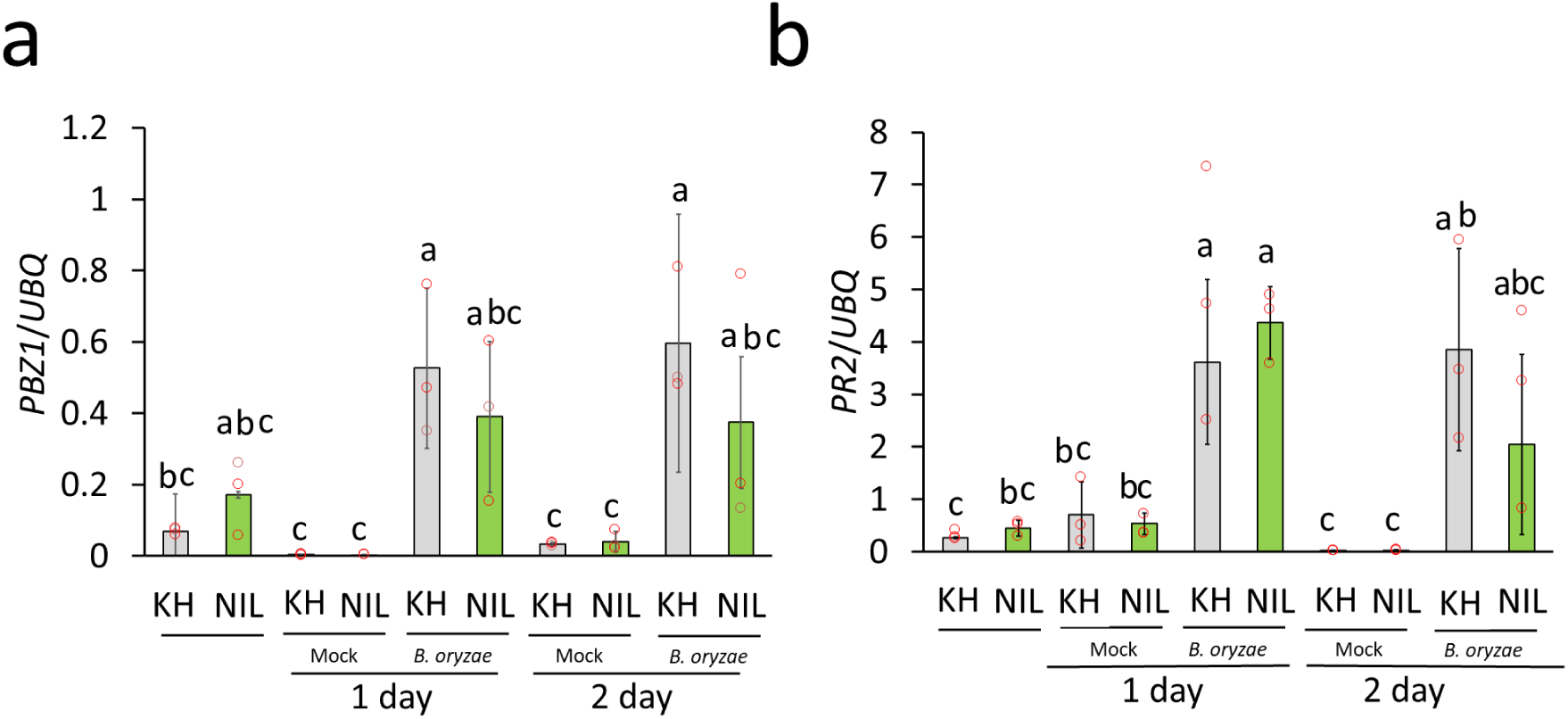
Expression of the pathogenesis-related genes *PBZ1* and *PR2* in leaves of Koshihikari (KH) and *bsr1*-NIL (NIL) after inoculation with *Bipolaris oryzae*. Leaves were collected 1 or 2 days after mock (water) or *B. oryzae* inoculation. Expression levels of *PBZ1* and *PR2* were normalized to that of ubiquitin. Data are presented as mean ± s.d.; n = 3. Different letters indicate significant differences (α = 0.05, Tukey’s HSD test).

