## Supporting Information for "Rice brown spot resistance gene *bsr1* also confers resistance to bacterial blight by suppressing sucrose efflux"

**This PDF file includes:**

Figures S1 to S9
Tables S1 to S5
SI Reference

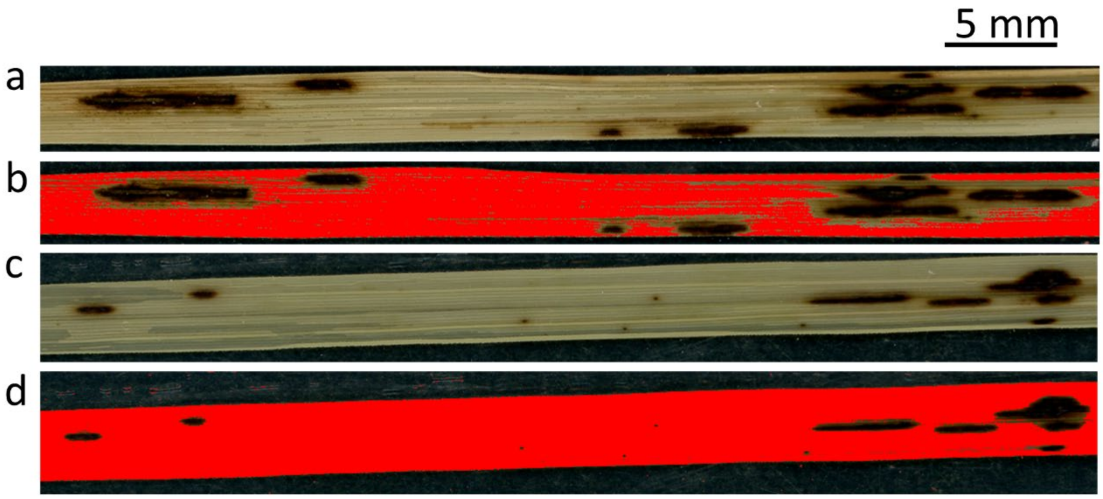


**Supplementary Fig. 1**

**Quantification of diseased leaf area. a,c,** Representative leaf images of Koshihikari **(a)** and *bsr1*-NIL **(c)**. Seedlings were inoculated with *Bipolaris oryzae*. Fourteen days after inoculation, leaves were detached and immersed in 70% ethanol to remove chlorophyll before imaging. **b,d,** Transformed images of Koshihikari **(b)** and *bsr1*-NIL **(d)** produced by ImageJ. Each pixel was classified as either diseased (black) or healthy (red). The percentage of lesion area was calculated as the ratio of diseased area to total leaf area. The diseased area was estimated as 36.8% in susceptible Koshihikari **(a, b)** and 4.2% in resistant *bsr1*-NIL **(c, d)**.

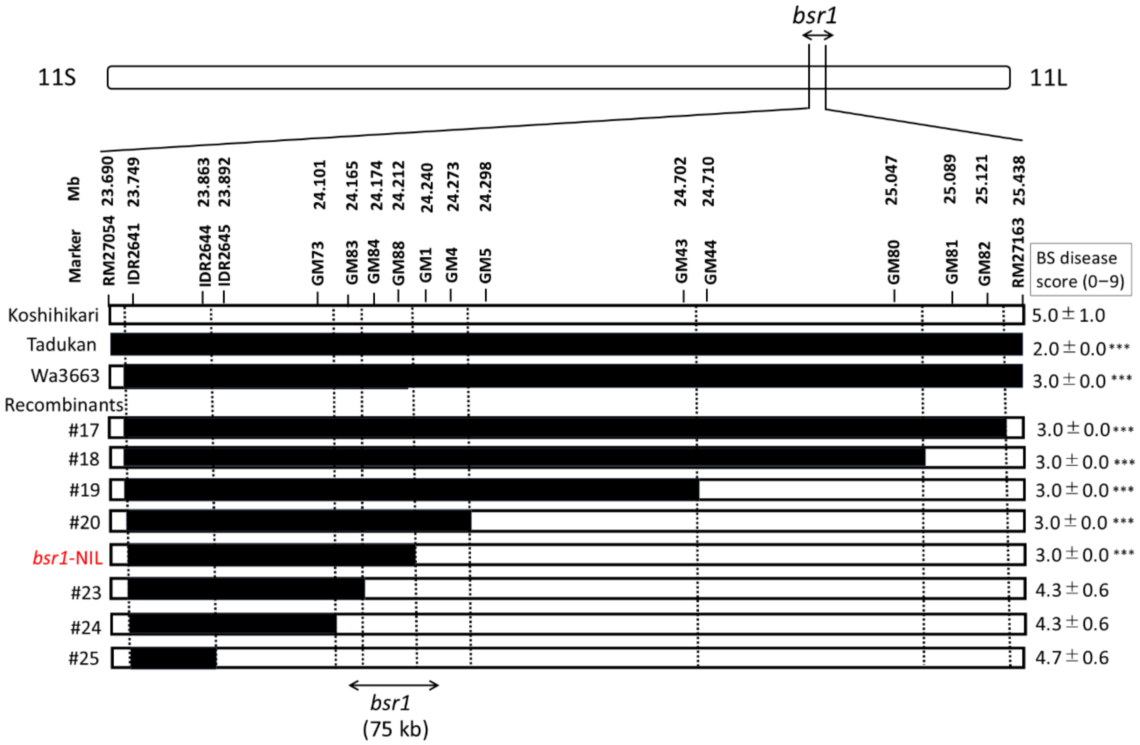


**Supplementary Fig. 2**

**Fine mapping of *bsr1* on the long arm of chromosome 11.** Black and white rectangles indicate chromosomal regions derived from Tadukan (brown spot [BS]-resistant) and Koshihikari (susceptible), respectively. Wa3663 is a BS-resistant line in which *bsr1* has been introduced into the Koshihikari genetic background; only the *bsr1* locus is derived from Tadukan[21,26]. Recombinants were identified among the progeny of the Wa3663 × Koshihikari cross. Marker positions are based on the International Rice Genome Sequencing Project 1.0 pseudomolecules of the Nipponbare genome. The candidate quantitative trait locus (*bsr1*), indicated at the bottom of the figure, was defined based on phenotypic data obtained from by BS resistance field tests. BS disease scores (means ± s.d.; 11 plants × 3 repeats) are shown to the right of the recombinant lines. Asterisks indicate significant differences compared with Koshihikari (Dunnett's test, $***P < 0.001$).

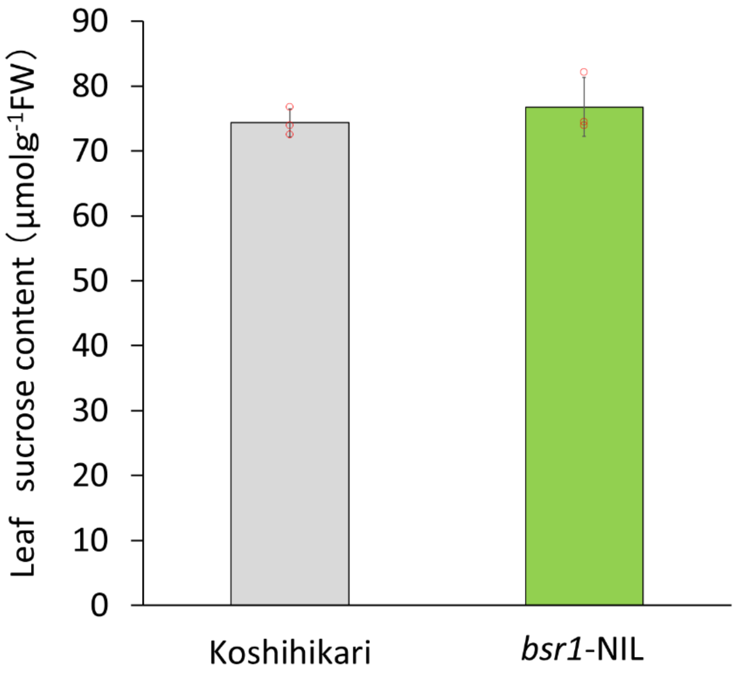


**Supplementary Fig. 3**

**Sucrose content of leaf blades at the ripening stage in Koshihikari and *bsr1*-NIL.**

Leaves were taken 105 days after sowing. Data are presented as mean ± s.d.; n = 3.

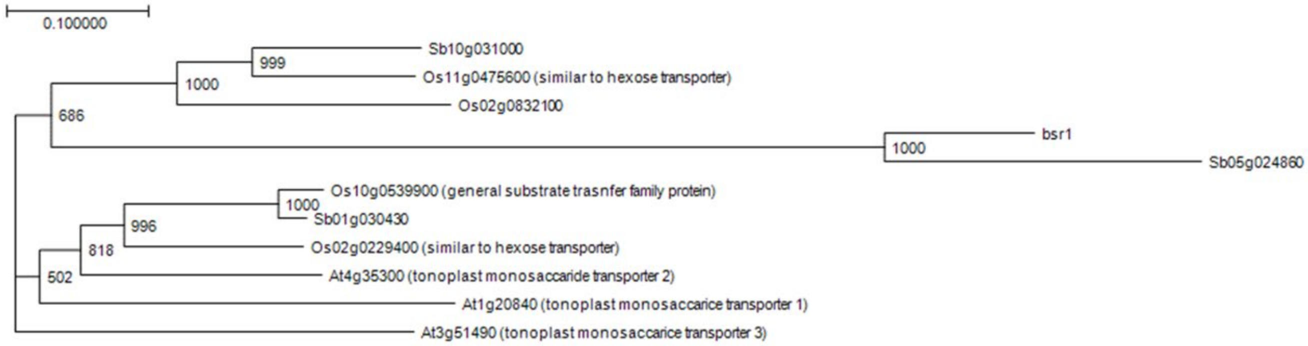


**Supplementary Fig. 4**

**Phylogenetic analysis of *bsr1*.** The phylogenic tree shows the predicted *bsr1* homologs encoding a conserved sucrose transporter domain in *Oryza sativa* (*Os*), *Sorghum bicolor* (*Sb*), and *Arabidopsis thaliana* (*At*). Homologous proteins were identified by BLAST searches using the BSR1 amino acid sequence as the query. Sequences were aligned using Genetyx ver. 16, and a phylogenetic tree was constructed using the neighbor-joining method. The number at each node represents the bootstrap value out of a total of 1000 bootstraps.

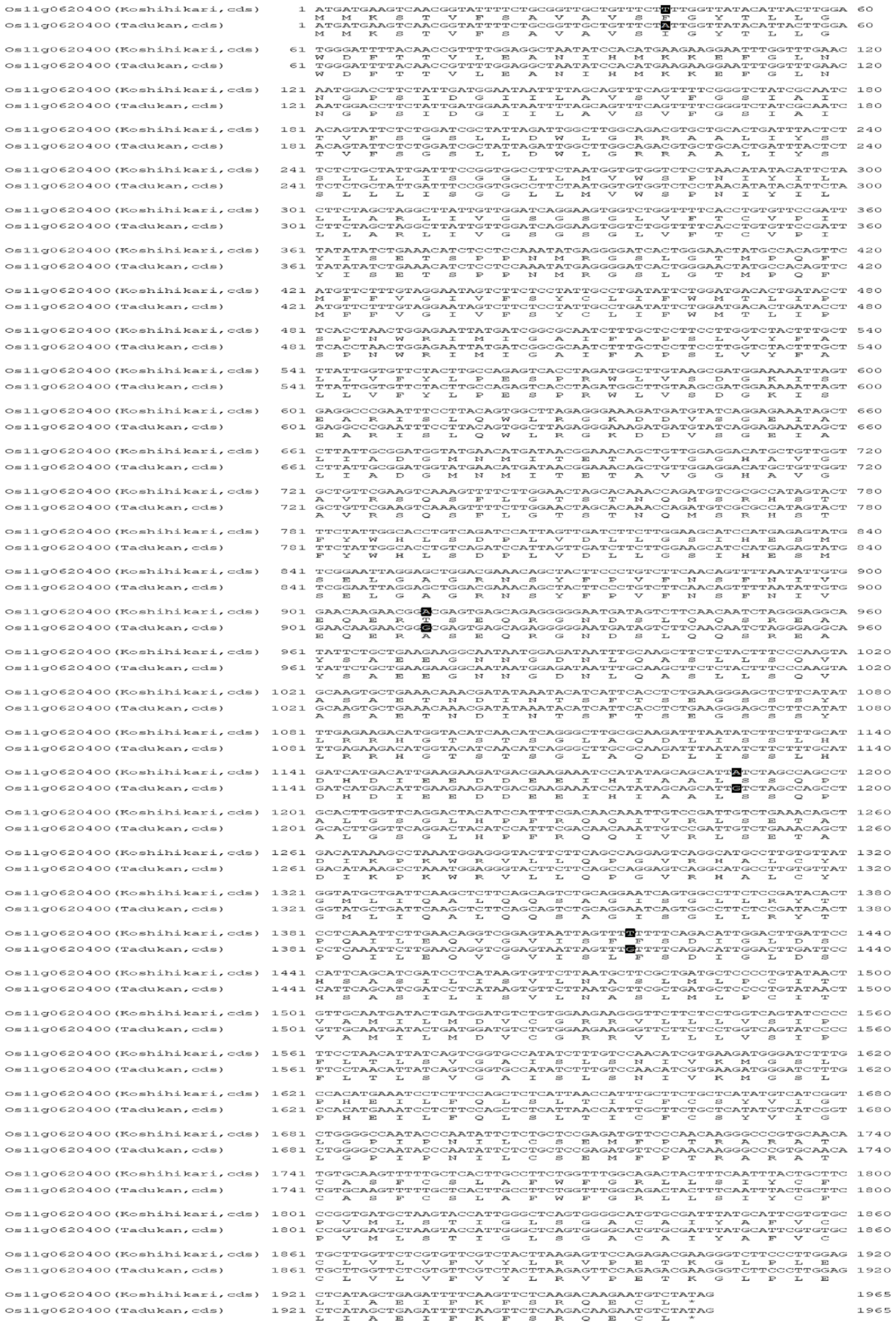

**Supplementary Fig. 5**

**Amino acid sequences of the *bsr1* (*Os11g0620400*) product in Koshihikari and Tadukan.** The four black boxes indicate single-nucleotide polymorphisms between Koshihikari and Tadukan.

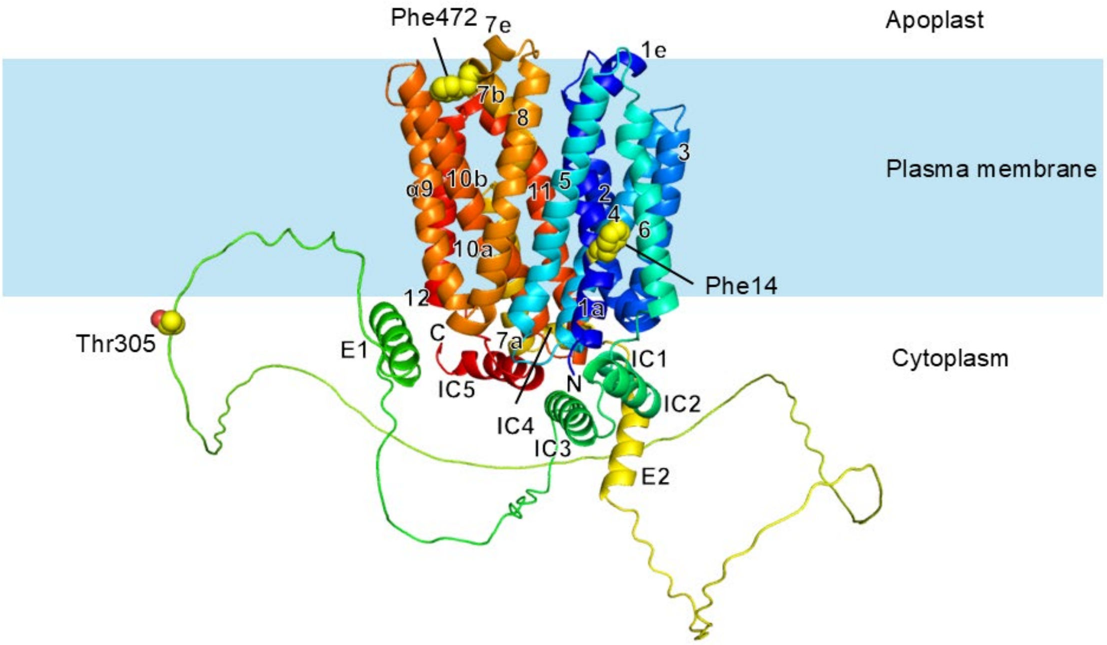


**Supplementary Fig. 6**

**AlphaFold-predicted structure of BSR1.** The main part of the protein consists of a transmembrane domain formed by 12 transmembrane helices (1–12). The intracellular face of the domain is attached to an intracellular helical domain, which is composed of five intracellular helices (IC1–5). These overall folds are identical to those of the glucose transporter GLUT31[1]. There is an extra-long loop region spanning approximately 200 residues and containing at least two additional helices (E1 and E2) between helices IC3 and IC4. The side chains of the three non-conserved residues between Koshihikari and Tadukan are shown as yellow spheres. Phe14 and Phe472 are found in the transmembrane domain, whereas Thr305 is located in the intracellular long loop region.

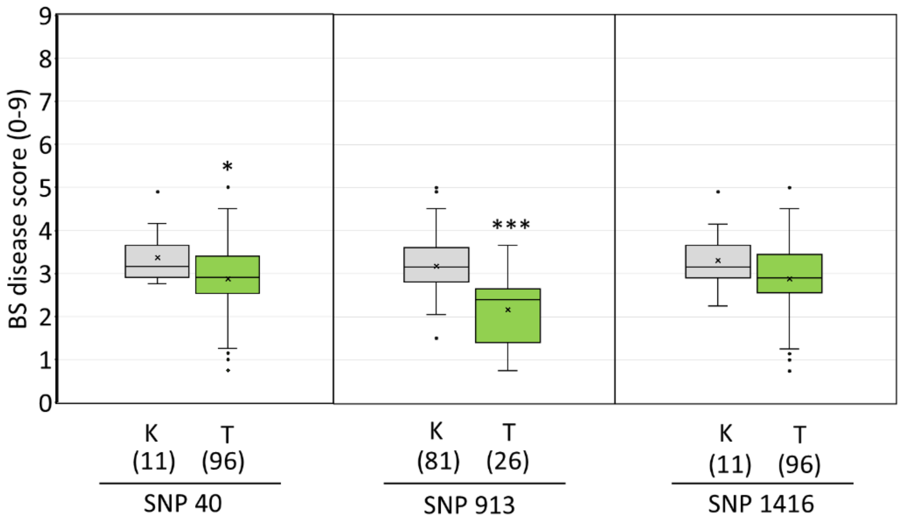


**Supplementary Fig. 7**

**Associations between contributions of *bsr1* SNP genotypes and brown spot resistance in cultivars from the World Rice Collection and Japanese Rice Collection listed in Supplementary Table 2.** SNP40, SNP913, and SNP1416 result in amino acid substitutions between the Koshihikari and Tadukan *bsr1* alleles. Box plots show the effect of each SNP on brown spot resistance. K: Koshihikari-type SNP; T: Tadukan-type SNP. The number of cultivars with each genotype is shown in parentheses. Asterisks indicate significant differences compared with Koshihikari (Student's *t*-test, *$P < 0.05$, ***$P < 0.001$).

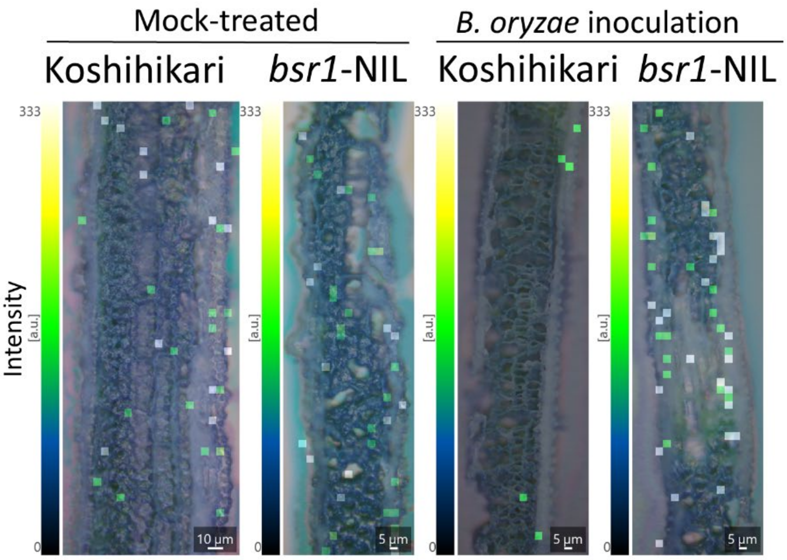


**Supplementary Fig. 8**

**Visualization of sucrose (*m*/*z* 381.08) in *Bipolaris oryzae*-inoculated leaves of Koshihikari and *bsr1*-NIL.** Twenty-two-day-old plants were inoculated with *B. oryzae*, and segments from the middle portion of fully expanded fourth leaf blades were embedded in optimum cutting temperature compound three days after inoculation. Frozen samples were sectioned longitudinally (parallel to the leaf vein) at a thickness of 10 µm using a cryostat and mounted onto indium tin oxide–coated glass slides for mass spectrometry imaging (MSI) analysis of sucrose distribution. For MSI analysis of leaf cross-sections, modified laser irradiation conditions, including a reduced laser spot size (approx. 5 µm), were used to achieve higher spatial resolution. The right side of each image corresponds to the leaf surface.

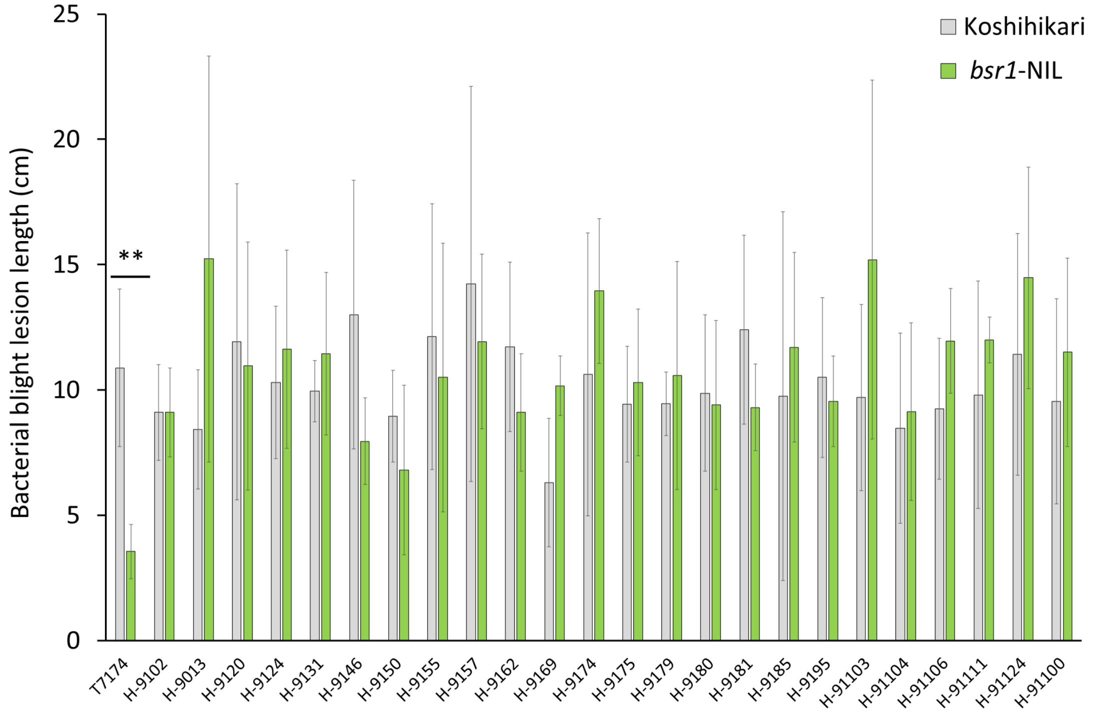


**Supplementary Fig. 9**

**Evaluation of bacterial blight resistance of Koshihikari and *bsr1*-NIL.** Flag leaves were inoculated with race I *Xanthomonas oryzae* pv. *oryzae* strains using the clipping method, and lesion lengths were measured 14 days after inoculation. Data are presented as mean ± s.d. of 3 or 4 biological replicates. Asterisks indicate significant differences compared with Koshihikari (Student's *t*-test, $^{**}P < 0.01$).

Supplementary Table 1 Predicted genes mapped in the 31.2-kb genomic region of the *bsr1* locus according to the Nipponbare reference genome

| Gene number | ID | Location | Annotation |
|---|---|---|---|
| g1 | Os11g0620400 (*bsr1* ) | 24184610–24191330 | Sugar/inositol transporter domain containing protein<br>Monosaccharide transporter (AZT subfamily), AZT protein 6 |
| g2 | Os11g0620450 | 24187723–24190926 | Hypothetical protein |
| g3 | Os11g0620500 | 24192139–24195237 | Tyrosine protein kinase domain containing protein |
| g4 | Os11g0620800 | 24203624–24204382 | Hypothetical conserved gene |
| g5 | LOC_Os11g40564.1 | 24209515–24208418 | Expressed protein (no gene ontology annotation found) |
| g6 | Os11g0621000 | 24210656–24214593 | Protein of unknown function DUF716 family protein |

Supplementary Table 2 Sequence polymorphisms in the exon region of *bsr1* and brown spot (BS) field resistance among World Rice Collection (WRC) and Japan Rice Collection (JRC) cultivars.

| Accession no. | Cultivar name | Origin | Subgroup | SNPs detected between Tadukan and Koshihikari in the exon region of *bsr1* | | | | BS disease score by field test (Matsumoto *et al.* *Breeding Research*, 2017) | | | |
|---|---|---|---|---|---|---|---|---|---|---|---|
| | | | | Exon 1 | Exon 3 | Exon 3 | Exon 4 | Year | | Average | SD |
| | | | | A/T | G/A[a] | G/A | G/T | | | | |
| | | | | Ile/Phe | Ala/Thr | S.S.[b] | Leu/Phe | 2014 | 2015 | | |
| | | | | 40[c] | 913[c] | 1188[c] | 1416[c] | | | | |
| | Koshihikari (susceptible) | Japan | *Temperate japonica* | T | A | A | T | 4.7 | 3.6 | 4.15 | 0.78 |
| WRC 20 | Tadukan (resistant, donor of *bsr1* ) | Philippines | *Indica* | A | G | G | G | 1.3 | 1.5 | 1.40 | 0.14 |
| WRC 05 | Naba | India | *Indica* | A | G | G | G | 2.8 | 2.5 | 2.65 | 0.21 |
| WRC 06 | Puluik Arang | Indonesia | *Indica* | A | G | G | G | 2.5 | 2.8 | 2.65 | 0.21 |
| WRC 07 | Davao 1 | Philippines | *Indica* | A | G | G | G | 2.3 | 2.5 | 2.40 | 0.14 |
| WRC 09 | Ryou Suisan Koumai | China | *Indica* | A | G | G | G | 3.5 | 2.5 | 3.00 | 0.71 |
| WRC 11 | Jinguoyin | China | *Indica* | A | G | G | G | 4.5 | 2.8 | 3.65 | 1.20 |
| WRC 12 | Dahonggu | China | *Indica* | A | G | G | G | 1.3 | 1.5 | 1.40 | 0.14 |
| WRC 13 | Asu | China | *Indica* | A | G | G | G | 2.8 | 2.5 | 2.65 | 0.21 |
| WRC 14 | IR 58 | Philippines | *Indica* | A | G | G | G | 0.5 | 1.0 | 0.75 | 0.35 |
| WRC 15 | Co 13 | India | *Indica* | A | G | G | G | 2.8 | 2.8 | 2.80 | 0.00 |
| WRC 17 | Keiboba | China | *Indica* | A | G | G | G | 2.5 | 2.8 | 2.65 | 0.21 |
| WRC 18 | Qingyu (Seiyu) | Taiwan | *Indica* | A | G | G | G | 1.8 | 2.0 | 1.90 | 0.14 |
| WRC 22 | Calotoc | Philippines | *Indica* | A | G | G | G | 1.0 | 1.0 | 1.00 | 0.00 |
| WRC 24 | Pinulupot 1 | Philippines | *Indica* | A | G | G | G | 1.8 | 1.0 | 1.40 | 0.57 |
| WRC 27 | Nepal 8 | Nepal | *Indica* | A | G | G | G | 3.3 | 2.8 | 3.05 | 0.35 |
| WRC 44 | Basilanon | Philippines | *Indica* | A | G | G | G | 2.8 | 1.8 | 2.30 | 0.71 |
| WRC 45 | Ma sho | Myanmar | *Tropical japonica* | A | G | G | G | 2.8 | 2.0 | 2.40 | 0.57 |
| WRC 46 | Khao Nok | Laos | *Tropical japonica* | A | G | G | G | 2.0 | 1.5 | 1.75 | 0.35 |
| WRC 48 | Khau Mac Kho | Vietnam | *Tropical japonica* | A | G | G | G | 2.5 | 2.0 | 2.25 | 0.35 |
| WRC 52 | Khau Tan Chiem | Vietnam | *Tropical japonica* | A | G | G | G | 1.5 | 1.0 | 1.25 | 0.35 |
| WRC 57 | Milyang 23 | Korea | *Indica* | A | G | G | G | 2.5 | 2.5 | 2.50 | 0.00 |
| WRC 59 | Neang Phtong | Cambodia | *Indica* | A | G | G | G | - | - | - | - |
| WRC 60 | Hakphaynhay | Laos | *Indica* | A | G | G | G | - | - | - | - |
| WRC 61 | Radin Goi Sesat | Malaysia | *Indica* | A | G | G | G | - | - | - | - |
| WRC 63 | Bleiyo | Thailand | *Indica* | A | G | G | G | - | - | - | - |
| WRC 65 | Rambhog | Indonesia | *Indica* | A | G | G | G | - | - | - | - |
| WRC 66 | Bingala | Myanmar | *Indica* | A | G | G | G | - | - | - | - |
| WRC 68 | Khao Nam Jen | Laos | *Temperate japonica* | A | G | G | G | - | - | - | - |
| WRC 97 | Chin Galay | Myanmar | *Indica* | A | G | G | G | - | - | - | - |
| WRC 98 | Deejiaohualuo | China | *Indica* | A | G | G | G | 2.0 | 2.0 | 2.00 | 0.00 |
| WRC 99 | Hong Cheuh Zai | China | *Indica* | A | G | G | G | 2.3 | 2.8 | 2.55 | 0.35 |
| WRC 100 | Vandaran | Sri Lanka | *Indica* | A | G | G | G | 2.8 | 2.5 | 2.65 | 0.21 |
| JRC 50 | Himenomochi | Japan | *Temperate japonica* | A | G | G | G | 2.5 | 2.3 | 2.40 | 0.14 |
| WRC 53 | Tima | Bhutan | *Tropical japonica* | A | G | A | G | 1.3 | 1.0 | 1.15 | 0.21 |
| WRC 16 | Vary Futsi | Madagascar | *Indica* | A | A | G | G | 2.8 | 2.5 | 2.65 | 0.21 |
| WRC 21 | Shwe Nang Gyi | Myanmar | *Indica* | A | A | G | G | 1.0 | 2.0 | 1.50 | 0.71 |
| WRC 25 | Muha | India | *Indica* | A | A | G | G | 4.0 | 3.5 | 3.75 | 0.35 |
| WRC 26 | Jhona 2 | India | *Indica* | A | A | G | G | 3.5 | 3.0 | 3.25 | 0.35 |
| WRC 01 | Nipponbare | Japan | *Temperate japonica* | A | A | A | G | 4.0 | 3.3 | 3.65 | 0.49 |
| WRC 02 | Kasalath | India | *Indica* | A | A | A | G | 4.8 | 3.8 | 4.30 | 0.71 |
| WRC 03 | Bei Khe | Cambodia | *Indica* | A | A | A | G | 3.0 | 3.0 | 3.00 | 0.00 |
| WRC 04 | Jena 035 | Nepal | *Indica* | A | A | A | G | 2.3 | 2.8 | 2.55 | 0.35 |
| WRC 10 | Qiu Zhao Zong | China | *Indica* | A | A | A | G | 4.0 | 3.0 | 3.50 | 0.71 |
| WRC 19 | Deng Pao Zhai | China | *Indica* | A | A | A | G | 2.8 | 2.5 | 2.65 | 0.21 |
| WRC 23 | Lebed | Philippines | *Indica* | A | A | A | G | 2.5 | 2.0 | 2.25 | 0.35 |
| WRC 28 | Jarjan | Bhutan | *Indica* | A | A | A | G | 2.5 | 2.8 | 2.65 | 0.21 |
| WRC 29 | Kalo Dhan | Nepal | *Indica* | A | A | A | G | 3.3 | 3.0 | 3.15 | 0.21 |
| WRC 30 | Anjana Dhan | Nepal | *Indica* | A | A | A | G | 3.0 | 2.8 | 2.90 | 0.14 |
| WRC 31 | Shoni | Bangladesh | *Indica* | A | A | A | G | 4.0 | 3.0 | 3.50 | 0.71 |
| WRC 32 | Tupa 121-3 | Bangladesh | *Indica* | A | A | A | G | 3.0 | 3.0 | 3.00 | 0.00 |
| WRC 33 | Surjamukhi | India | *Indica* | A | A | A | G | 3.3 | 3.0 | 3.15 | 0.21 |
| WRC 34 | ARC 7291 | India | *Indica* | A | A | A | G | 4.0 | 3.5 | 3.75 | 0.35 |
| WRC 35 | ARC 5955 | India | *Indica* | A | A | A | G | 4.0 | 3.5 | 3.75 | 0.35 |
| WRC 36 | Ratul | India | *Indica* | A | A | A | G | 3.0 | 2.5 | 2.75 | 0.35 |
| WRC 37 | ARC 7047 | India | *Indica* | A | A | A | G | 4.0 | 3.5 | 3.75 | 0.35 |
| WRC 38 | ARC 11094 | India | *Indica* | A | A | A | G | - | - | - | - |
| WRC 39 | Badari Dhan | Nepal | *Indica* | A | A | A | G | 3.0 | 2.8 | 2.90 | 0.14 |
| WRC 40 | Nepal 555 | India | *Indica* | A | A | A | G | 2.8 | 2.3 | 2.55 | 0.35 |
| WRC 41 | Kaluheenati | Sri Lanka | *Indica* | A | A | A | G | 3.5 | 3.5 | 3.50 | 0.00 |
| WRC 42 | Local Basmati | India | *Indica* | A | A | A | G | 2.8 | 3.0 | 2.90 | 0.14 |
| WRC 43 | Dianyu 1 | China | *Temperate japonica* | A | A | A | G | 3.0 | 3.0 | 3.00 | 0.00 |
| WRC 49 | Padi Perak | Indonesia | *Tropical japonica* | A | A | A | G | 2.5 | 1.8 | 2.15 | 0.49 |
| WRC 50 | Rexmont | USA | *Tropical japonica* | A | A | A | G | 2.3 | 2.8 | 2.55 | 0.35 |
| WRC 51 | Urasan 1 | Japan | *Tropical japonica* | A | A | A | G | 5.5 | 3.5 | 4.50 | 1.41 |
| WRC 55 | Tupa 729 | Bangladesh | *Tropical japonica* | A | A | A | G | 3.0 | 2.8 | 2.90 | 0.14 |
| WRC 58 | Neang Menh | Cambodia | *Indica* | A | A | A | G | - | - | - | - |
| WRC 62 | Kemasin | Malaysia | *Indica* | A | A | A | G | - | - | - | - |
| WRC 64 | Padi Kuning | Indonesia | *Indica* | A | A | A | G | - | - | - | - |
| WRC 67 | Phulba | India | *Temperate japonica* | A | A | A | G | - | - | - | - |
| JRC 01 | Gaisen Mochi | Japan | *Tropical Japonica* | A | A | A | G | 2.8 | 2.8 | 2.80 | 0.00 |
| JRC 03 | Hinode | Japan | *Tropical Japonica* | A | A | A | G | 4.3 | 3.0 | 3.65 | 0.92 |
| JRC 04 | Senshou | Japan | *Tropical Japonica* | A | A | A | G | 4.0 | 3.3 | 3.65 | 0.49 |
| JRC 05 | Yamada Bake | Japan | *Tropical Japonica* | A | A | A | G | 3.3 | 2.8 | 3.05 | 0.35 |
| JRC 06 | Kaneko B | Japan | *Tropical Japonica* | A | A | A | G | 3.3 | 3.0 | 3.15 | 0.21 |
| JRC 07 | Iruma Nishiki | Japan | *Tropical Japonica* | A | A | A | G | 4.3 | 3.0 | 3.65 | 0.92 |
| JRC 08 | Okka Modoshi | Japan | *Tropical Japonica* | A | A | A | G | 3.3 | 3.0 | 3.15 | 0.21 |
| JRC 10 | Hirayama | Japan | *Tropical Japonica* | A | A | A | G | 4.0 | 3.3 | 3.65 | 0.49 |
| JRC 11 | Kahei | Japan | *Tropical Japonica* | A | A | A | G | 3.8 | 3.3 | 3.55 | 0.35 |
| JRC 12 | Oiran | Japan | *Tropical Japonica* | A | A | A | G | 3.8 | 3.3 | 3.55 | 0.35 |
| JRC 13 | Bouzu Mochi | Japan | *Tropical Japonica* | A | A | A | G | 3.5 | 3.0 | 3.25 | 0.35 |

| | | | | | | | | | | | |
|---|---|---|---|---|---|---|---|---|---|---|---|
| JRC 14 | Meguro Mochi | Japan | *Tropical Japonica* | A | A | A | G | 3.3 | 2.8 | 3.05 | 0.35 |
| JRC 17 | Akage | Japan | *Temperate japonica* | A | A | A | G | 5.5 | 4.5 | 5.00 | 0.71 |
| JRC 18 | Hassokuho | Japan | *Temperate japonica* | A | A | A | G | 2.5 | 2.5 | 2.50 | 0.00 |
| JRC 19 | Wataribune | Japan | *Temperate japonica* | A | A | A | G | 3.8 | 2.5 | 3.15 | 0.92 |
| JRC 20 | Hosogara | Japan | *Temperate japonica* | A | A | A | G | 4.3 | 3.3 | 3.80 | 0.71 |
| JRC 21 | Akamai | Japan | *Temperate japonica* | A | A | A | G | 3.3 | 3.3 | 3.30 | 0.00 |
| JRC 22 | Mansaku | Japan | *Temperate japonica* | A | A | A | G | 3.8 | 3.5 | 3.65 | 0.21 |
| JRC 24 | Joushuu | Japan | *Temperate japonica* | A | A | A | G | 3.3 | 3.3 | 3.30 | 0.00 |
| JRC 28 | Shinriki Mochi | Japan | *Temperate japonica* | A | A | A | G | 3.3 | 2.8 | 3.05 | 0.35 |
| JRC 29 | Shichimenchou Mochi | Japan | *Temperate japonica* | A | A | A | G | 3.0 | 2.8 | 2.90 | 0.14 |
| JRC 31 | Kameji | Japan | *Temperate japonica* | A | A | A | G | 2.5 | 2.5 | 2.50 | 0.00 |
| JRC 32 | Omachi | Japan | *Temperate japonica* | A | A | A | G | 2.5 | 2.5 | 2.50 | 0.00 |
| JRC 33 | Shinriki | Japan | *Temperate japonica* | A | A | A | G | 2.8 | 2.8 | 2.80 | 0.00 |
| JRC 35 | Kabashiko | Japan | *Temperate japonica* | A | A | A | G | 3.3 | 3.0 | 3.15 | 0.21 |
| JRC 36 | Sekiyama | Japan | *Temperate japonica* | A | A | A | G | 4.0 | 3.5 | 3.75 | 0.35 |
| JRC 37 | Shinyamadaho 2 | Japan | *Temperate japonica* | A | A | A | G | 2.5 | 2.0 | 2.25 | 0.35 |
| JRC 38 | Nagoya Shiro | Japan | *Temperate japonica* | A | A | A | G | 3.3 | 3.3 | 3.30 | 0.00 |
| JRC 39 | Shiroine (Kemomi) | Japan | *Temperate japonica* | A | A | A | G | 3.3 | 2.8 | 3.05 | 0.35 |
| JRC 41 | Akamai | Japan | *Indica* | A | A | A | G | 3.0 | 3.0 | 3.00 | 0.00 |
| JRC 42 | Touboshi | Japan | *Indica* | A | A | A | G | 2.8 | 2.8 | 2.80 | 0.00 |
| JRC 43 | Akamai | Japan | *Indica* | A | A | A | G | 3.0 | 3.0 | 3.00 | 0.00 |
| JRC 44 | Karahoushi | Japan | *Indica* | A | A | A | G | 3.5 | 3.0 | 3.25 | 0.35 |
| JRC 45 | Hiyadachitou | Japan | *Temperate japonica* | A | A | A | G | 4.0 | 3.8 | 3.90 | 0.14 |
| JRC 47 | Okabo | Japan | *Temperate japonica* | A | A | A | G | 2.8 | 2.5 | 2.65 | 0.21 |
| JRC 48 | Hakamuri (Yokoyama) | Japan | *Temperate japonica* | A | A | A | G | 3.0 | 2.8 | 2.90 | 0.14 |
| JRC 49 | Rikutou Rikuu 2 | Japan | *Temperate japonica* | A | A | A | G | 4.3 | 3.3 | 3.80 | 0.71 |
| JRC 51 | Shinshuu | Japan | *Temperate japonica* | A | A | A | G | 3.5 | 3.5 | 3.50 | 0.00 |
| JRC 53 | Raiden | Japan | *Temperate japonica* | A | A | A | G | 2.3 | 1.8 | 2.05 | 0.35 |
| JRC 54 | Houmaishinden Ine | Japan | *Temperate japonica* | A | A | A | G | 3.3 | 2.0 | 2.65 | 0.92 |
| WRC 47 | Jaguary | Brazil | *Tropical japonica* | T | A | A | G | 3.8 | 2.0 | 2.90 | 1.27 |
| JRC 23 | Ishijiro | Japan | *Temperate japonica* | T | A | A | T | 3.0 | 2.5 | 2.75 | 0.35 |
| JRC 25 | Dango | Japan | *Temperate japonica* | T | A | A | T | 3.5 | 2.3 | 2.90 | 0.85 |
| JRC 26 | Aikoku | Japan | *Temperate japonica* | T | A | A | T | 3.5 | 3.3 | 3.40 | 0.14 |
| JRC 27 | Ginbouzu | Japan | *Temperate japonica* | T | A | A | T | 3.5 | 2.8 | 3.15 | 0.49 |
| JRC 30 | Morita Wase | Japan | *Temperate japonica* | T | A | A | T | 4.0 | 3.3 | 3.65 | 0.49 |
| JRC 34 | Kyotoasahi | Japan | *Temperate japonica* | T | A | A | T | 3.3 | 3.0 | 3.15 | 0.21 |
| JRC 40 | Akamai | Japan | *Indica* | T | A | A | T | 3.3 | 2.5 | 2.90 | 0.57 |
| JRC 46 | Fukoku | Japan | *Temperate japonica* | T | A | A | T | 6.0 | 3.8 | 4.90 | 1.56 |
| JRC 52 | Aichiasahi | Japan | *Temperate japonica* | T | A | A | T | 3.3 | 3.0 | 3.15 | 0.21 |

Shaded boxes indicate the Tadukan-type SNPs.

[a] SNPs infered to be associated with *bsr1* -mediated resistance.

[b] Synonymous substitution.

[c] SNP position relative to the start of the coding sequence.

Supplementary Table 3 Sequence polymorphisms of *bsr1* exon 3 among *Oryza rufipogon* accessions[a]

| Accession no. | Origin | G/A<br>Ala/Thr<br>913[b] | SNP type[c] |
|---|---|---|---|
| W1294 | Philippines | G | Tadukan |
| W1981 | Indonesia | G | Tadukan |
| W2051 | Bangladesh | G | Tadukan |
| IRGC105715 | -[d] | G | Tadukan |
| W0106 | India | A | Koshihikari |
| W0120 | India | A | Koshihikari |
| W0137 | India | A | Koshihikari |
| W0180 | Thailand | A | Koshihikari |
| W0593 | Malaysia | A | Koshihikari |
| W0630 | Burma | A | Koshihikari |
| W1230 | Indonesia | A | Koshihikari |
| W1236 | Papua New Guinea | A | Koshihikari |
| W1551 | Thailand | A | Koshihikari |
| W1669 | India | A | Koshihikari |
| W1681 | India | A | Koshihikari |
| W1715 | China | A | Koshihikari |
| W1807 | Sri Lanka | A | Koshihikari |
| W1866 | Thailand | A | Koshihikari |
| W1886 | Thailand | A | Koshihikari |
| W1921 | Thailand | A | Koshihikari |
| W1945 | -[d] | A | Koshihikari |
| W1962 | China | A | Koshihikari |
| W1965 | China | A | Koshihikari |
| W1976 | Indonesia | A | Koshihikari |
| W2003 | India | A | Koshihikari |
| W2057 | Bangladesh | A | Koshihikari |
| W2078 | Australia | A | Koshihikari |
| W2109 | Australia | A | Koshihikari |
| W2114 | Australia | A | Koshihikari |
| W2117 | Australia | A | Koshihikari |
| W2263 | Cambodia | A | Koshihikari |
| IRGC105444 | -[d] | A | Koshihikari |
| IRGC105908 | -[d] | A | Koshihikari |

[a] Data were obtained from OryzaGenome2.1 (http://viewer.shigen.info/oryzagenome21detail/index.xhtml).

[b] SNP position relative to the start of the coding sequence.

[c] Accessions were categorized according to the nucleotide at SNP913 (position 913 from the start of the coding sequence).

[d] Origin unknown.

Supplementary Table 4 Fungal and bacterial strains used in this study.

| **MAFF no.** | **Name** | **Scientific name** | **Collection area in Japan** | **Race** |
|---|---|---|---|---|
| 235499 | T.AOKI AR0126 | *Bipolaris oryzae* | Okinawa (24.34°N, 124.18°E) | - |
| 305066 | 13 | *Bipolaris oryzae* | Hokkaido (43.06°N, 141.35°E) | - |
| 305067 | F-1 | *Bipolaris oryzae* | Ehime (33.83°N, 132.76°E) | - |
| 305197 | 445 | *Bipolaris oryzae* | Osaka (34.69°N, 135.50°E) | - |
| 305453 | 1297 | *Bipolaris oryzae* | Miyazaki (31.91°N, 131.42°E) | - |
| 510752 | - | *Bipolaris oryzae* | Chiba (35.83°N, 140.26°E) | - |
| 311018 | T7174 | *Xanthomonas oryzae* pv. *oryzae* | Kyoto (35.01°N, 135.77°E) | I |
| 311019 | T7147 | *Xanthomonas oryzae* pv. *oryzae* | Gifu (35.42°N, 136.76°E) | II |
| 311020 | T7133 | *Xanthomonas oryzae* pv. *oryzae* | Mie (33.81°N, 136.05°E) | III |
| 301235 | T7177 | *Xanthomonas oryzae* pv. *oryzae* | Kyoto (35.30°N, 135.26°E) | I |
| 210549 | H-9102 | *Xanthomonas oryzae* pv. *oryzae* | Ehime (33.28°N, 132.73°E) | I |
| 210550 | H-9013 | *Xanthomonas oryzae* pv. *oryzae* | Ishikawa (36.56°N, 136.65°E) | I |
| 210567 | H-9120 | *Xanthomonas oryzae* pv. *oryzae* | Niigata (37.92°N, 139.07°E) | I |
| 210571 | H-9124 | *Xanthomonas oryzae* pv. *oryzae* | Toyama (36.70°N, 137.21°E) | I |
| 210578 | H-9131 | *Xanthomonas oryzae* pv. *oryzae* | Ishikawa (36.67°N, 136.73°E) | I |
| 210593 | H-9146 | *Xanthomonas oryzae* pv. *oryzae* | Miyagi (38.00°N, 140.62°E) | I |
| 210597 | H-9150 | *Xanthomonas oryzae* pv. *oryzae* | Kumamoto (32.85°N, 131.14°E) | I |
| 210602 | H-9155 | *Xanthomonas oryzae* pv. *oryzae* | Aichi (34.90°N, 137.50°E) | I |
| 210604 | H-9157 | *Xanthomonas oryzae* pv. *oryzae* | Ishikawa (36.33°N, 136.53°E) | I |
| 210609 | H-9162 | *Xanthomonas oryzae* pv. *oryzae* | Akita (40.21°N, 140.03°E) | I |
| 210616 | H-9169 | *Xanthomonas oryzae* pv. *oryzae* | Shimane (35.37°N, 132.75°E) | I |
| 210621 | H-9174 | *Xanthomonas oryzae* pv. *oryzae* | Tokushima (33.92°N, 134.66°E) | I |
| 210622 | H-9175 | *Xanthomonas oryzae* pv. *oryzae* | Ishikawa (36.33°N, 136.53°E) | I |
| 210626 | H-9179 | *Xanthomonas oryzae* pv. *oryzae* | Kochi (32.94°N, 132.73°E) | I |
| 210627 | H-9180 | *Xanthomonas oryzae* pv. *oryzae* | Shiga (34.97°N, 136.17°E) | I |
| 210628 | H-9181 | *Xanthomonas oryzae* pv. *oryzae* | Fukui (36.06°N, 136.22°E) | I |
| 210632 | H-9185 | *Xanthomonas oryzae* pv. *oryzae* | Oita (33.10°N, 131.41°E) | I |
| 210642 | H-9195 | *Xanthomonas oryzae* pv. *oryzae* | Yamagata (38.73°N, 139.83°E) | I |
| 210650 | H-91103 | *Xanthomonas oryzae* pv. *oryzae* | Yamaguchi (34.53°N, 131.58°E) | I |
| 210651 | H-91104 | *Xanthomonas oryzae* pv. *oryzae* | Hyogo (34.89°N, 135.23°E) | I |
| 210653 | H-91106 | *Xanthomonas oryzae* pv. *oryzae* | Shiga (35.33°N, 136.06°E) | I |
| 210658 | H-91111 | *Xanthomonas oryzae* pv. *oryzae* | Hiroshima (35.50°N, 136.78°E) | I |
| 210671 | H-91124 | *Xanthomonas oryzae* pv. *oryzae* | Yamaguchi (34.18°N, 131.47°E) | I |
| 210647 | H-91100 | *Xanthomonas oryzae* pv. *oryzae* | Tochigi (37.01°N, 136.78°E) | I |

Supplementary Table 5 Primers used in this study

| **Name** | **Forward primer (5′–3′)** | **Reverse primer (5′–3′)** | **Purpose** | **Position** | **Genotype (Tadukan/Koshihikari)** |
|---|---|---|---|---|---|
| RM27054 | CGGCATGGGTTATAGCAAGACC | GGAAGATGCGACAATTGACATGG | Mapping of *bsr1* (SSR marker) | | |
| RM27163 | TCATCTCGTAAAATTTCGATTCCGATG | TAATCACCCGGTGCAACGCA | Mapping of *bsr1* (SSR marker) | | |
| IDR2641 | CGCATCTCAACCGTCTTCTG | TTATAGCACGCCGAGGTCAT | Mapping of *bsr1* (InDel marker) | | |
| IDR2644 | GCCCGTTTCCTAAGGCTTTT | GTCCCTACCATAGCCAGTCC | Mapping of *bsr1* (InDel marker) | | |
| IDR2645 | CCGAGGGATTTTGCACACAT | TCGATGACAGTCCAAGGATCA | Mapping of *bsr1* (InDel marker) | | |
| GM1 | TGGGTCCTACGTGTCAATGA | AGCCCAATTAAGCTGCCCTA | Mapping of *bsr1* (InDel marker) | | |
| GM4 | ATATGTCGTGTCTTGTGGAATTTT | TCCACCGAATTGATTTTCGCT | Mapping of *bsr1* (InDel marker) | | |
| GM5 | GACCAGTGATTTAGCCGACG | GTGTGTTTGGTTCCACGTCA | Mapping of *bsr1* (InDel marker) | | |
| GM43 | AGAGAAACCAGAGACAAGGTAAA | TGGAGGGATCTCATGCAGTT | Mapping of *bsr1* (InDel marker) | | |
| GM44 | CAACCGGGAGTAGCAATTCG | CCTAGTCCCGTTGTTGTTACC | Mapping of *bsr1* (InDel marker) | | |
| GM73 | AACTGGTGAATCGATGGGCT | CCAGCTAATTGCCACAACGA | Mapping of *bsr1* (InDel marker) | | |
| GM80 | CCACGATCTGGGCTTCAAGA | GCCGAAAATGATGAAAATGGCA | Mapping of *bsr1* (InDel marker) | | |
| GM81 | GTCACGTGGCATCAGTTACG | CGTGCACATCAAGAACGCTA | Mapping of *bsr1* (InDel marker) | | |
| GM82 | GAGCTTTTGAACTGGCGCTA | TTTTGGCCCAAAACAAAATTGG | Mapping of *bsr1* (InDel marker) | | |
| GM83 | TCTTAAGGCTGTCATGGCGA | GTTCCCAATTTAGGCAGCCA | Mapping of *bsr1* (InDel marker) | | |
| GM84 | AGCTCGATCCAGATCAGTGG | GTCACCTGTTTTGCCCTGTT | Mapping of *bsr1* (InDel marker) | | |
| GM86 | TGTGTACAGGGACAGATGGT | CTCAGACAACCGTTCACGAA | Mapping of *bsr1* (InDel marker) | | |
| GM88 | GTCACGTGGCATCAGTTACG | CGTGCACATCAAGAACGCTA | Mapping of *bsr1* (InDel marker) | | |
| GM91 | GGAGCTTGACGCTTTTTGAC | AAAATACACTGGTTATGTGCCATTT | Mapping of *bsr1* (Sequence analysis) | 24228667 | T/G |
| GM92 | ACCCTTTGCGTGAGGAGTAA | CCTAATCTGTTGGAGTGGAGGA | Mapping of *bsr1* (Sequence analysis) | 24215460 | C/T |
| GM95 | CCCATTCATTTAAGTGGAAACAA | ATGGGAGACAAGGCTACGTG | Mapping of *bsr1* (Sequence analysis) | 24225868 | G/C |
| GM99 | TGCCGGTTCCTACTGGTTAG | TGGACTGTAGCCAACACATACC | Mapping of *bsr1* (Sequence analysis) | 24179329-24179330 | AG/GA |
| GM100 | GGATGGTTTTCCTTTTTAGCC | TTAAAGTGCTACTCAGGTCATTGC | Mapping of *bsr1* (Sequence analysis) | 24183607 | C/T |
| GM101 | AATCTTAGGCGGCGTTCAAT | AAAAACGCAAGCTGGTTGAC | Mapping of *bsr1* (Sequence analysis) | 24185049 | A/G |
| GM105 | ACCGTCCGATATGGATCAAA | GGTTGTGTGTTAGCCGCAAT | Mapping of *bsr1* (Sequence analysis) | 24214848 | A/G |
| GM106 | AGAGGCAAATGGCTTCTGAG | CACCACCAATCCACTCCTCT | Mapping of *bsr1* (Sequence analysis) | 24213246 | T/G |
| CRISPR-bsr1 | gttgAACCAAATTCCTTCTTCATG | aaacCATGAAGAAGGAATTTGGTT | *bsr1* CRISPR-Cas guide RNA | | |
| GM134 | CCATGTTGTACGTCAAAAGAAGA | GATTGCGATAGACCCGAAAA | Sequence analysis of CRISPR-Cas transformants | | |
| bsr1-ORF-U | ATCCGGTACCGAATTATGATGAAGTCAACGGTATTTTCTG | | Subcellular localization | | |
| bsr1-ORF-L | GTCTAGATATCTCGATAGACATTCTTGTCTTGAGAACTTG | | Subcellular localization | | |
| bsr1 | GCCCGAATTTCCTTACAGTG | GCGACATCTGGTTTGTGCTA | qRT-PCR for *bsr1* (SYBR) | | |
| PR2 | AAGATTGTTCTGAGAAGAGATCGATCGA | GCTACGCGAAAATAGGTCTGGTAAACTT | qRT-PCR for *PR2* | | |
| PR2-T | ATGGCGTTGCTTCCGTTTTAACACTGG | | TaqMan for *PR2* (FAM) | | |
| PBZ1 | CAAATTCTCGTGGCGTTTGA | AAGGCACATAAACACAACCACAA | qRT-PCR for *PBZ1* | | |
| PBZ1-T | CCGTGAGAGTAATTTCGTGCTTTAAG | | TaqMan for *PBZ1* (FAM) | | |
| Ubiquitin | GAGCCTCTGTTCGTCAAGTA | ACTCGATGGTCCATTAAACC | qRT-PCR for *Ubiquitin* (SYBR, TaqMan) | | |
| Ubiquitin-T | TTGTGGTGCTGATGTCTACTTGTGTC | | TaqMan for *Ubiquitin* (VIC) | | |
| Os11g0620400 (*bsr1*) | GGCCCGAATTTCCTTACAGT | GCGACATCTGGTTTGTGCTA | RT-PCR | | |
| Os11g0620450 | GCAAAGATCCCATCTTCACG | CCCTTAATGCTTCGCTGATG | RT-PCR | | |
| Os11g0620500 | CCATTACCGCAAGGAAGAAA | AGCTTCTCGTGGAGGTACGA | RT-PCR | | |
| Os11g0620800 | GTTCCGCGAGGAGTTAATTG | AACGCAGCCACAAAAGAGAT | RT-PCR | | |
| Os11g0621000 | GGCATGCTCCTCATGTTCTT | GACGACGACGAGGAGGTAGT | RT-PCR | | |
| LOC_Os11g40564.1 | GGGATGGAGCATCATCTGG | CATCCCACTTGGTATTGACG | RT-PCR | | |
